# HB-EGF and zinc activate EGFR to induce reactive neural stem cells in the mouse hippocampus after seizures

**DOI:** 10.1101/2022.11.02.514820

**Authors:** Oier Pastor-Alonso, Irene Durá, Sara Bernardo-Castro, Emilio Varea, Teresa Muro-García, Soraya Martín-Suárez, Juan Manuel Encinas-Pérez, Jose Ramon Pineda

**Affiliations:** Laboratory of Neural Stem Cells and Neurogenesis, Achucarro Basque Center for Neuroscience. Scientific Park, 48940 Leioa, Bizkaia, Spain; Faculty of Biology. University of Valencia. 46100 Valencia, Spain; Ikerbasque, The Basque Foundation for Science. Euskadi Plaza, 5, 48009 Bilbao, Bizkaia, Spain; Department of Neurosciences, University of the Basque Country (UPV/EHU). Scientific Park, 48940 Leioa, Bizkaia, Spain; Signaling lab. Department of Cell Biology and Histology. Faculty of Medicine and Nursing, University of the Basque Country (UPV/EHU), Leioa, Bizkaia, Spain

**Keywords:** Neural stem cells, gliosis, hippocampus, epilepsy, epidermal growth factor receptor, zinc

## Abstract

Hippocampal seizures mimicking mesial temporal lobe epilepsy (MTLE) cause a profound disruption of the adult neurogenic niche in mice. Seizures provoke neural stem cells to switch to a reactive phenotype (reactive-neural stem cells, React-NSCs)) characterized by multibranched hypertrophic morphology, massive activation to enter mitosis, symmetric division and final differentiation into reactive astrocytes. As a result, neurogenesis is chronically impaired. Here we, using a mouse model of MTLE, show that the epidermal growth factor receptor (EGFR) signalization pathway is key for the induction of React-NSCs and that its inhibition exerts a beneficial effect on the neurogenic niche. We show that during the initial days after the induction of seizures by a single intrahippocampal injection of kainic acid, a strong release of zinc and heparin-binding epidermal growth factor, both activators of the EGFR signalization pathway in neural stem cells, is produced. Administration of the EGFR inhibitor gefitinib, a chemotherapeutic in clinical phase IV, prevents the induction of React-NSCs and preserves neurogenesis.

**Significance:** In mouse models of MTLE-HS, seizures cause a profound disruption of the hippocampal neurogenic niche and neurogenesis results chronically impaired, in agreement with what occurs in the human MTLE-HS hippocampus. Thus, the normal cognitive functions associated with neurogenesis are altered, but also the endogenous regenerative capacity that could compensate the high rate of neurons in the granule cell layer of the dentate gyrus. We provide here for the first time a molecular mechanism (the EGFR transduction pathway) regulating the induction of React-NSCs.

## INTRODUCTION

Mesial temporal lobe epilepsy (MTLE) is the most common form of epilepsy in adults (Falconer et al., 1964; Margerison and Corsellis, 1966). A portion of patients develop MTLE with hippocampal sclerosis (MTLE-HS) consisting in unilateral hippocampal atrophy, neuronal death, reactive gliosis and granule cell dispersion (GCD). MTLE-HS associates with drug-resistance often leading to unilateral hippocampalectomy as a last-resort therapeutic strategy (Crespel et al., 2005; Tatum, 2012). In the dentate gyrus (DG) of the hippocampus of most mammals, possibly including humans (Eriksson et al., 1998; Spalding et al., 2013; Llorens-Bobadilla et al., 2015), a population of neural stem cells (NSCs) continues to generate new neurons postnatally and throughout adulthood (Encinas and Enikolopov, 2008; Spalding et al., 2013). In human MTLE-HS as well as in experimental models based on the intrahippocampal injection of kainic acid (KA), reduction or absence of markers of neurogenesis and cell proliferation in the hippocampal neurogenic niche have been reported (Heinrich et al., 2006; Ledergerber et al., 2006; Nitta et al., 2008). In other models of MTLE (based on pilocarpine administration or the electrical induction of seizures) neurogenesis was found to be increased (Parent et al., 1997; Madsen et al., 2000). In parallel to alterations in neurogenesis, seizures trigger astrogliogenesis in the DG. We have recently characterized using mouse models of MTLE-HS, how seizures induce the transformation of hippocampal NSCs into reactive NSCs (React-NSCs) who contribute to HS through the direct generation of reactive astrocytes at the expense of neurogenesis (Sierra et al., 2015a; Muro-García et al., 2019; Valcárcel-Martín et al., 2020).

The mechanisms controlling the differentiation of NSCs into React-NSCs in the intrahippocampal KA mouse model have not yet been investigated and could be of therapeutic use as preserving the normal population of hippocampal NSCs in MTLE-HS could: a) preserve neurogenesis and its normal functions; 2) allow for the endogenous restoration of the GCL neurons dead by excitotoxicity and 3) reduce reactive gliosis. We herein studied the involvement of the epidermal growth factor receptor (EGFR) in the induction of React-NSCs and whether it could be a potential target to preserve them after seizures. EGFR has been directly reported to be present in proliferative embryonic neural precursors (Ciccolini et al., 2005), as well as in activated hippocampal adult NSCs and amplifying neural progenitors (Jhaveri et al., 2015; Walker et al., 2016). Although in normal conditions brain astroglia expresses weakly EGFR it plays a role in the proliferation and differentiation of astrocytes (Simpson et al., 1982). Also, a dramatic increase of EGFR has been observed after brain injury or focal ischemia, especially in reactive astrocytes and microglia (Nieto-Sampedro et al., 1988; Planas et al., 1998). Increased levels of EGFR promote the adult NSCs ability to differentiate into astrocytes (Burrows et al., 1997) and asymmetric distribution of EGFR in late cortical progenitors promotes astrocyte lineage in cells expressing high levels of EGFR (Sun et al., 2005). Moreover, it has been recently postulated that the genetic deletion of EGFR affects the self-renewal capacity of neural stem and progenitor cells (NSPCs) from the subventricular zone (SVZ), promoting the differentiation of the progeny towards astrocytes (Robson et al., 2018). However, mice lacking EGFR show reduced numbers of cortical astrocytes and increased hypersensitivity to epileptic seizures induced by intraperitoneal KA administration (Sibilia et al., 2007; Robson et al., 2018). Further, our previous data on functional gene analysis from the hippocampus of mice subjected to MTLE-HS showed, among others, an early upregulation of the ErbB transduction pathway (Sierra et al., 2015b). The ErbB protein family contains four structurally related receptor tyrosine kinases, including the EGFR. Both EGFR and fibroblastic growth factor receptor (FGFR) signaling pathways have been demonstrated to be highly mitogenic for astrocytes (Simpson et al., 1982; Lewis et al., 1992). Also, FGFR expression increased in the cortex and hippocampus of rats injected intraperitoneally with KA (Van Der Wal et al., 1994), but no information exists regarding EGFR.

## RESULTS

### Seizures cause an early overexpression and activation of the EGFR signaling pathway in the hippocampus

We first sought to evaluate the expression of both EGFR and FGFR in a standardized model of MTLE-HS consisting in a single administration of 1 nmol of KA in the DG of c57bl/6 mice (Bouilleret et al., 1999; Sierra et al., 2015b). The hippocampus was collected at early time points after the KA injection (1.5h, 12h, 24h, and 72h) (Figure 1A) and analyzed by real-time quantitative polymerase chain reaction (RT-qPCR) (Figure 1B). A 2-3-fold significant increase in EGFR mRNA was found as early as 1.5h and was maintained at 24h and 72h post-KA (One-way ANOVA p=0.028, p=0.037 and p=0.007 respectively compared to controls; Figure 1B). In order to measure the activation of the EGFR pathway we determined the phosphorylation of the receptor (P-EGFR) on its tyrosine site 845 (Tyr845), which has been shown to be required for triggering DNA synthesis (Boerner et al., 2005). The western blotting (WB) analysis revealed a progressive increase of P-Tyr845 (respect to the normal EGFR) that became significant at 24h and 72h post-KA (3.116±0.698 at 24h and 8.540±0.357 at 72h compared to 1.000±0.385 control; One-way ANOVA p=0.04 and p<0.001 respectively; Figure 1C and 1D). In addition, we measured the activation level, also by phosphorylation, of different downstream effectors of the EGFR signaling pathway in the hippocampal samples (Figure 1C). EGFR mediates the activation of Janus kinase/signal transducer and activator of transcription 3 (STAT3) (Ueno et al., 1997); AKT, involved in cell survival; and ERK1/2, which regulates cell stress and proliferation (Meloche and Pouysségur, 2007; Lill and Sever, 2012; Goffin and Zbuk, 2013). Our results showed that STAT3 was strongly activated 12-fold compared to control at 12h post-KA injection and then decreased (Kruskal Wallis p=0.026; Figure 1E). AKT phosphorylation increased gradually, becoming significant at 72h (15-fold increase respect to control; Kruskal Wallis p=0.043; Figure 1F). Phosphorylation of ERK1/2 was more prominent, becoming statistically significant as early as 1.5h post-KA (8-fold respect to control) and then remaining elevated afterwards (10-20-fold respect to control; One-way ANOVA p=0.023, p=0.05, p=0.05 and p<0.001 respectively compared to control; Figure 1G). In contrast, no differences were found in the amount of FGFR1 or FGFR2 mRNA (Supplementary Figure 1A), or in FGFR1 protein (Supplementary Figure 1B), in the hippocampus of MTLE-HS mice compared to control ones.

**FIGURE 1.**
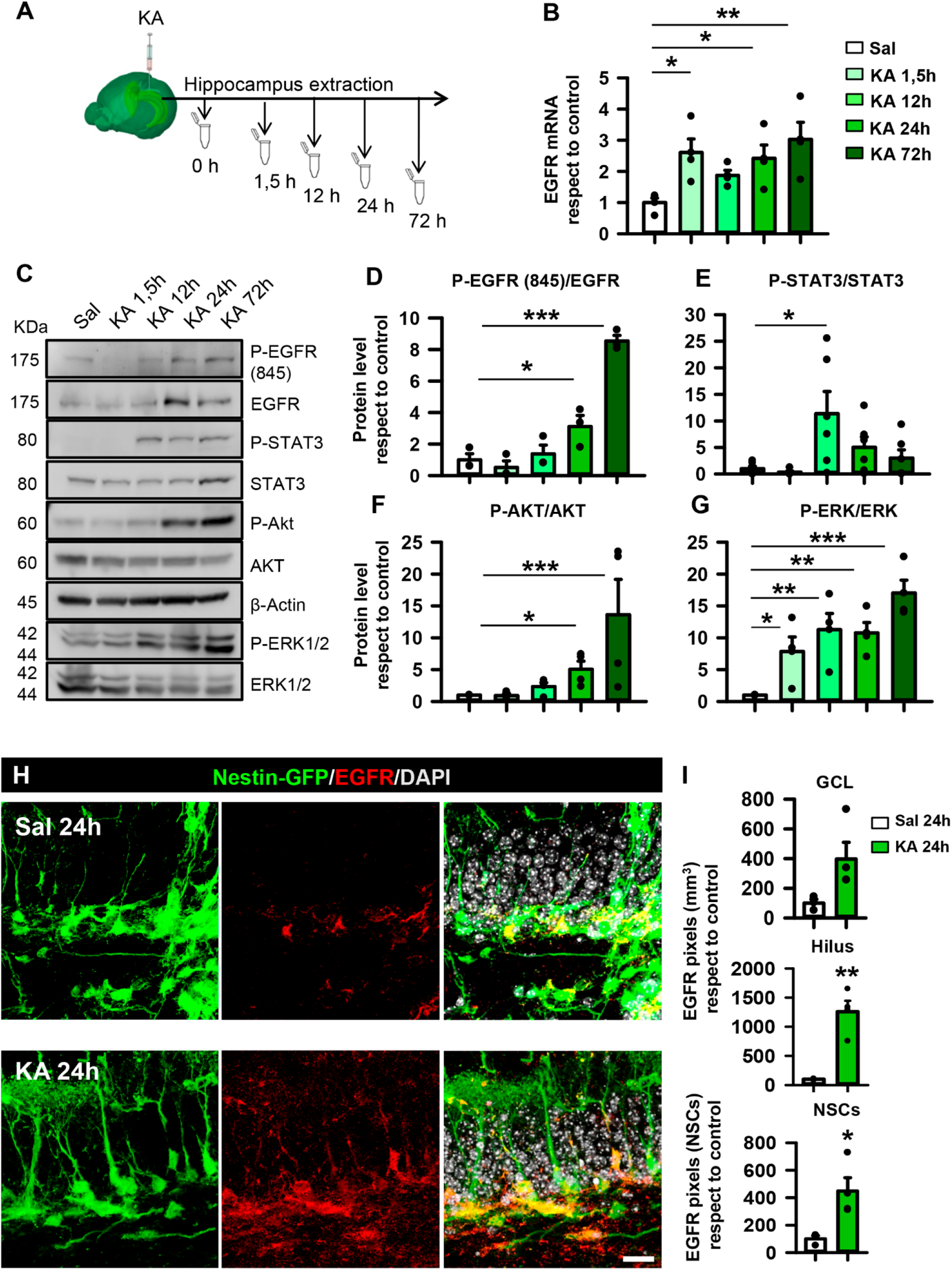
Increased expression of EGFR in the hippocampal neurogenic niche and activation of the EGFR signaling pathway are early events in MTLE-HS. A, Schematic time-course of hippocampus dissection after intrahippocampal injection of KA (1nmol). B, Determination of EGFR mRNA expression by RT-qPCR showing an early increase after KA injection. One-way ANOVA *p < 0.05, **p < 0.01. Bars show mean ± SEM. n=3. Dots show individual data. C, Determination of changes in normal and phosphorylated protein levels involved in EGFR signaling by WB of hippocampal samples. p-EGFR, p-STAT3, p-AKT and p-ERK are increased. Data are representative of 3 biological replicates, with representative blots cropped and marked with solid lines. D, Quantification of the ratio of phosphorylated versus non-phosphorylated forms of EGFR. E, Quantification of the ratio of phosphorylated versus non-phosphorylated forms of STAT3. F, Quantification of the ratio of phosphorylated versus non-phosphorylated forms of AKT. G, Quantification of the ratio of phosphorylated versus non-phosphorylated forms of ERK after WB. Phosphorylation changes are compared respect to control after WB. One-way ANOVA in (D, G) and Kruskal-Wallis in (E, F) *p < 0.05, **p < 0.01, ***p < 0.001. Bars show mean ± SEM. Dots show individual data. n=5. (H) Confocal microscopy images after immunostaining for EGFR in the SGZ of the hippocampus of control (upper panel) and MTLE-HS (lower panel) Nestin-GFP showing increased expression in NSCs and ANPs early after intrahippocampal KA injection. Scale bar 20 μm. I, Quantification of EGFR immunofluorescent signal in the GCL, hilar region and NSCs. Mann-Whitney for GCL and Student’s t-test for hilar region and NSCs *p < 0.05, **p < 0.01. Bars show mean ± SEM. Dots show individual data. n=3.

### EGFR expression increases after seizures in hippocampal NSPCs

EGFR plays a central role in promoting cell proliferation through the Ras/MAPK/ERK pathway (Schlessinger and Bar-Sagi, 1994; Cobb, 1999). Moreover, nuclear EGFR has been strongly correlated with high cell proliferation (Lin et al., 2001). Because a much higher rate of cell division is one of the React-NSCs characteristics, we hypothesize that EGFR expression would be increased in MTLE-HS-induced React-NSCs. Thus, we determined EGFR expression in the hippocampal neurogenic niche of control and MTLE-HS mice. We used 2-month-old (2-m.o.) transgenic Nestin-GFP (on c57bl/6 background) mice in which NSCs and React-NSCs can be readily visualized (Sierra et al., 2015b; Muro-García et al., 2019). The mice were subjected to the intrahippocampal injection or KA (or saline for controls) and sacrificed 24h later. In controls, EGFR expression was sparse and tightly restricted to the neurogenic niche, specifically labeling a limited number of Nestin-GFP+ cells located in the subgranular zone (SGZ) of the DG (Figure 1H). In contrast, EGFR immunostaining had increased in the MTLE-HS mice and was noticeably prominent in more Nestin-GFP+ cells (Figure 1H). The quantification showed that EGFR expression (measured as pixels of positive immunostaining normalized for the respective regions) tended to be significantly increased in the GCL after KA (increase of 400% vs control levels; Student’s t-test p=0.057; Figure 1I). Expression of EGFR was also significantly increased in the hilus of the MTLE-HS mice (1200% vs control levels; Student’s t-test p=0.003; Figure 1I). Specifically, colocalization of EGFR with React-NSCs (MTLE-HS group) was increased by 450% compared to colocalization with control NSCs (Student’s t-test p=0.031; Figure 1I). We found no changes in FGFR1 expression by immunofluorescence in the hippocampal neurogenic niche (Supplementary Figure 1C), confirming the result obtained by RT-qPCR and WB (Supplementary Figure 1A and 1B).

### Pharmacological blockage of EGFR signaling reduces NSPC proliferation *in vitro* and *in vivo*

We next aimed to directly modulate EGFR signaling in NSPCs without the interference of the profusion of external niche signals present *in vivo*. We isolated and enriched hippocampal NSPCs *in vitro* (adaptation of (Pineda et al., 2013)) from 2-m.o. Nestin-GFP mice and confirmed first that one week after primary culture and neurosphere generation EGFR was present in all cells (Supplementary Figure 1D). However, NSPCs undergoing mitosis, as identified by their chromosome condensation typical of prophase and metaphase, expressed significantly higher levels of EGFR, quantified by measuring signal intensity after immunostaining for EGFR (2.34±0.18 vs. 0.81±0.05 pixel/μm^2^; Mann Whitney p<0.001; Supplementary Figure 2A and 2B). We then determined the effect of the pharmacological inhibition of EGFR signaling using the reversible inhibitor gefitinib, which disrupts EGFR kinase activity by reversibly binding within the ATP-binding pocket of the EGFR protein, efficiently inhibiting all tyrosine phosphorylation sites on EGFR (Pedersen et al., 2005) (Figure 2A and 2B). We checked the effect of EGFR inhibition on both EGFR Tyr845 and Tyr1068, located within the activation loop of the kinase domain of EGFR. WB analyses showed that the addition of gefitinib (2 μM) for 1h into EGF-stimulated (20 ng/mL) cultured NSPCs efficiently reduced phosphorylation at Tyr845 by 81.65% and at Tyr1068 by 69.47% respect to control cultures (Mann Whitney p=0.029 for Tyr845 and Student’s t-test p=0.031 for Tyr1068; Figure 2A and 2B). Moreover, the presence of gefitinib efficiently reduced phosphorylation of the downstream effector ERK1/2 by 75.92% respect to controls (Student’s t-test p=0.012; Figure 2A and Figure 2B). To corroborate the effects on cell proliferation, we cultured NSPCs in presence of 2 μM gefitinib for 48h and a pulse of 10 μM 5-bromo-2’-deoxyuridine (BrdU) was administered 1h before fixation in order to identify dividing cells (Figure 2C). We then co-stained for DAPI, BrdU and Nestin-GFP (Figure 2D). Our results showed a 37% reduction of BrdU-positive cells in presence of gefitinib respect to the control (Mann-Whitney p<0.001; Figure 2E). We also tested the effect of inhibiting EGFR on NSPCs using afatinib, an irreversible inhibitor of EGFR. NSPCs that were cultured for 24h with afatinib (2 μM) had reduced proliferation passing from a 20.27±5.95% of them being positive for BrdU in controls to an 8.91±4.88% in afatinib-treated cells (Mann-Whitney p=0.046; Supplementary Figure 2C, 2D and 2E). However, this effect is attributable to the massive cell death induced by afatinib, as indicated by the dramatic loss of BrdU+ cells and Nestin-GFP+ NSPCs (plus the rounded and vacuolated morphology of the remaining ones) when NSPCs were cultured with afatinib for 48h (Mann Whitney p<0.001). As a confirmation of the deleterious effect of afatinib, the co-injection of KA plus afatinib (70 μM) in the hippocampus of Nestin-GFP mice also provoked a massive loss of Nestin-GFP+ and BrdU+ cells in the DG (Supplementary Figure 2H and 2I).

**FIGURE 2.**
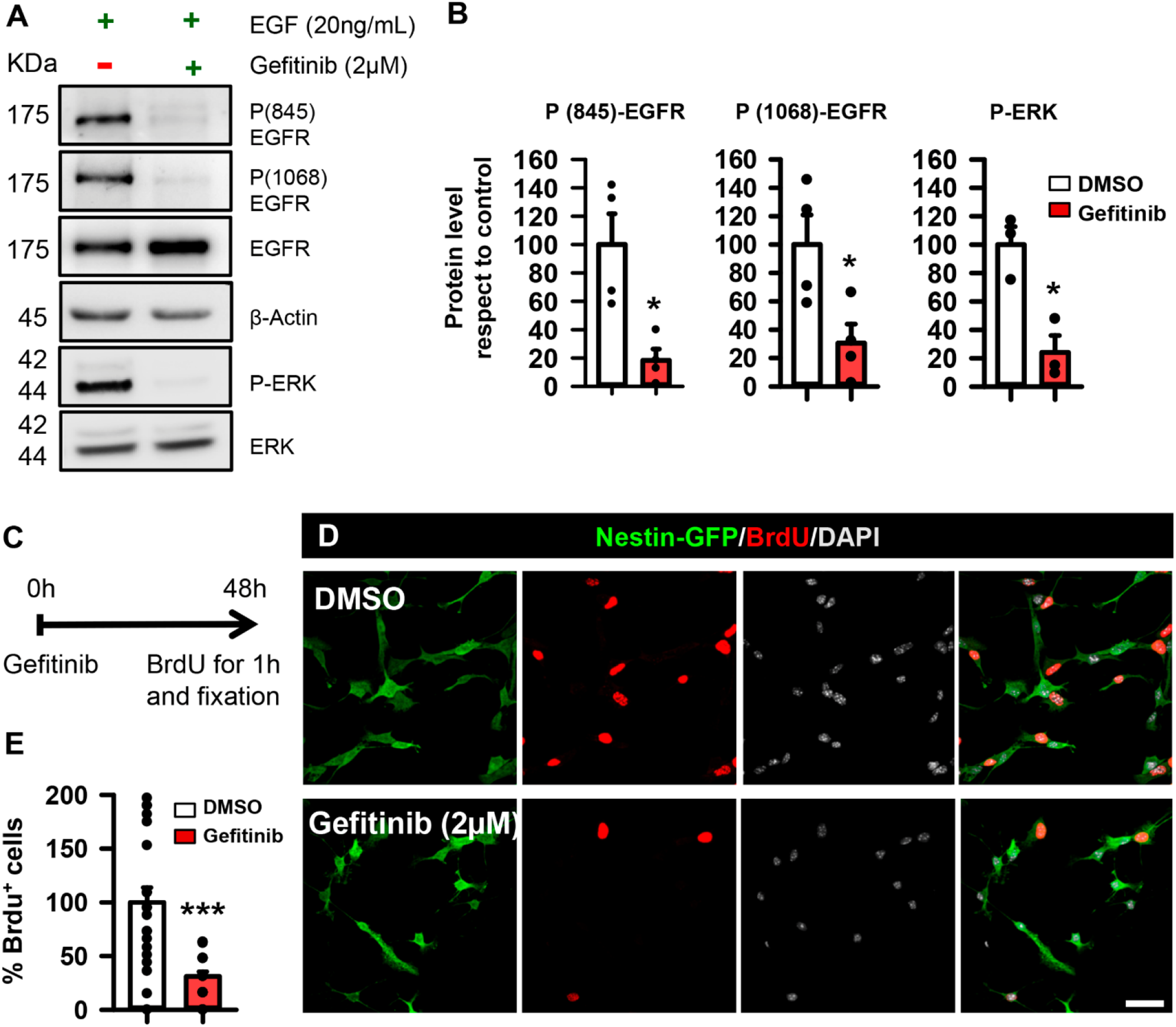
Inhibition of EGFR signaling with gefitinib reduces proliferation of NSPCs. A, EGFR of cultured hippocampal NSPCs stimulated with EGF was phosphorylated at Tyr845 and Tyr1068 residues as analyzed by WB. Its downstream effector ERK was also phosphorylated. Pretreatment with the EGFR inhibitor gefitinib blocked phosphorylation of the receptor, as well as of ERK. B, Quantification of the ratio of phosphorylated versus non-phosphorylated forms for EGFR Tyr845 and Tyr1068 and ERK (four independent replicates) showed the consistent inhibitory effect of gefitinib *p < 0.05, Mann Whitney for Tyr845 and Student’s t-test for Tyr1068 and ERK. Bars show mean ± SEM. Dots show individual data. n=4. C, Paradigm of the strategy to evaluate the effect on cultured cell proliferation. Cells were cultured in presence of EGF or gefitinib pretreatment plus EGF during 48h. A 1h-pulse of BrdU 10 μM was given before fixation to label mitotic cells. D, Representative immunofluorescence images of cultured NSPCs from Nestin-GFP mice showing the reduction of BrdU+ cells in gefitinib-treated cells. Scale bar 10 μm. E, Quantification of the number of BrdU+ cells expressed as percentage respect to total BrdU+ cells in the control condition showing the strong effect of gefitinib. Mann Whitney ***p < 0.001. Bars show mean ± SEM. Dots show random fields of two pooled independent experiments.

### Blocking EGFR signaling with gefitinib partially preserves hippocampal NSCs after MTLE-HS *in vivo*

We therefore chose to continue with gefitinib to study the effect of EGFR inhibition on the hippocampal neurogenic niche in the mouse model of MTLE-HS. As gefitinib does not cross the blood brain barrier, we administered it intranasally (McKillop et al., 2004), a pathway that has also been used for anti-epileptic drug treatment (Barakat et al., 2006). Although it has been reported that a single dose of gefitinib can block cell proliferation for 72h (Pedersen et al., 2005), we administered it twice a day starting right after KA injection to maximize its presence over the 3 days following the induction of MTLE-HS, when the peak of increase in proliferation is observed (Sierra et al., 2015b). The results showed a 57.25±4.98% reduction of the Ki67+ proliferating cells in the SGZ of the animals treated with gefitinib compared to the animals treated with the vehicle dimethyl sulfoxide (DMSO) (Student’s t-test p=0.002, Figure 3A and 3B). Likewise, the Ki67+ amplifying neural progenitors (ANPs) and NSCs also dropped to 32.02±5.25% and 45.25±5.92% respectively (Student’s t-test p=0.005 and p=0.007, Figure 3A and 3B).

**FIGURE 3.**
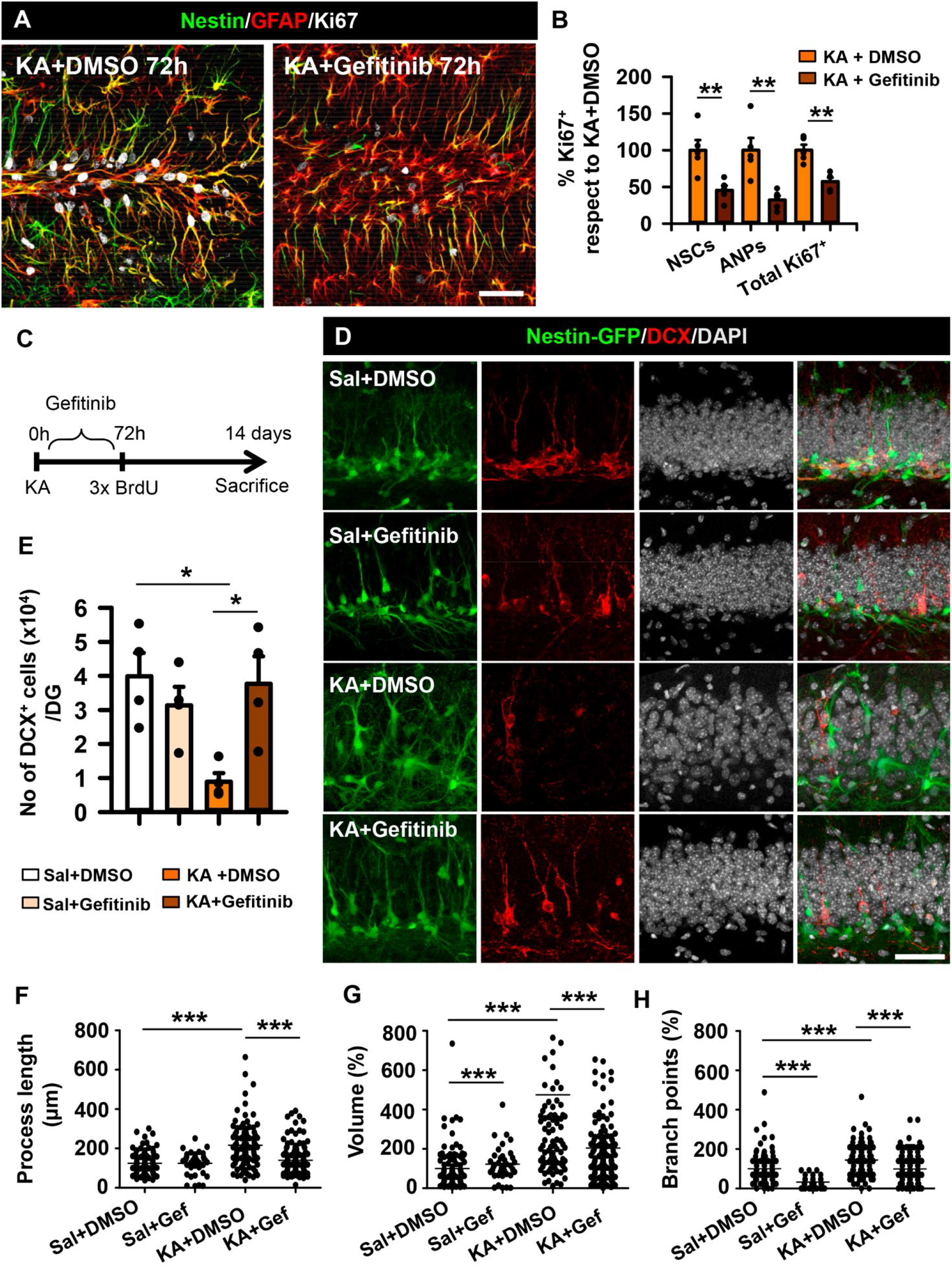
Inhibition of EGFR signaling with gefitinib in vivo reduces the induction of React-NSC induction and partially restores neurogenesis in MTLE-HS. A, Confocal microscopy images of MTLE-HS mice treated with intranasal gefitinib (10 mg/Kg) or vehicle, 3 days after intrahippocampal KA injection (1nmol) after staining for nestin, GFAP and Ki67. Scale bar, 50 μm. B, Quantification of proliferating (Ki67+) NSCs, ANPs and overall proliferation in the SGZ expressed as percentage respect to control. Student’s t-test **p < 0.01. Bars show mean ± SEM. Dots show individual data. n=4. C, Confocal microscopy images of DG sections after immunostaining for GFP, DAPI was used to label all cell nuclei. Scale bar, 20 μm. D, Quantification of changes in the length of the processes. E, Quantification of changes in cell volume. F, Quantification of changes in the number of branch points. Changes were quantified after 3D-sholl analysis and expressed as percentage respect to the control condition (sal+DMSO). One-way ANOVA ***p < 0.001. Dots show individual cells (D-F). G, Quantification of the volume of the GCL in saline+DMSO, saline+gefitinib, KA+DMSO and KA+gefitinib treated animals 14dpKA. Kruskal-Wallis *p < 0.05, **p < 0.01, ***p < 0.001. Bars show mean ± SEM. Dots show individual data. n=3.

In order to check the potential preservation of NSCs by EGFR inhibition in MTLE-HS we repeated the administration of gefitinib or DMSO in KA- or saline-injected Nestin-GFP mice but waited 14 days to sacrifice the mice and analyze them so that the transformation into React-NSCs is fully completed (Figure 3C and 3D).

In order to check the potential preservation of neurogenesis by EGFR inhibition in MTLE we repeated the administration of gefitinib or DMSO in KA- or saline-injected Nestin-GFP mice but waited 14 days to sacrifice and analyze them (Figure 3C and 3D). As expected, we found a severe depletion of DCX+ cells in the KA+DMSO mice (8914.61±2537.97 DCX+ cells) respect to saline+DMSO and saline+gefitinib (39888.21±6887.70 and 31341.6±5465.51 DCX+ cells respectively; One-way ANOVA p=0.022 and p=0.09; Figure 3D and 3E). In the MTLE mice treated with gefitinib, a substantial recovery of DCX+ cells was found in the DG, raising to 37641.67±8077.953 DCX+ cells (One-way ANOVA p=0.030; Figure 3D-E).

The reactive astrocyte-like morphology typical of the seizure-induced React-NSCs (Sierra et al., 2015b; Muro-García et al., 2019; Valcárcel-Martín et al., 2020) was reduced and NSCs looked closer to normal conserving their radial morphology in the MTLE-HS mice treated with gefitinib. To quantify this effect, we analyzed Nestin-GFP+ cells by Sholl analysis and measured the overall process length, the cell volume and ramification as branch points (Figure 3F, 3G and 3H, respectively). In the three parameters, gefitinib significantly reduced the effect of KA. Of notice, gefitinib also had an effect in saline-injected mice regarding cell volume and number of branch points. A pathological hallmark of MTLE-HS in the human hippocampus and animal models is granule cell dispersion (GCD). Which consists in the separation of granule cells resulting in increased length between the molecular layer and the hilus(Houser, 1990). We therefore tested whether GCD was also reversed by administration of gefitinib and found that indeed this was the case. An increase in GCD was found in the KA+DMSO animals (1.012±0.16 mm^3^; Kruskal Wallis p=0.007) respect to saline+DMSO (0.42±0.05 mm^3^; Kruskal Wallis p=0.01) and saline+gefitinib (0.36±0.04 mm^3^) and returned to normal values in the KA+gefitinib (0.46±0.02 mm^3^; Kruskal-Wallis p<0.001, Figure 3A and 3B).

Finally, we analyzed the effect of gefitinib in MTLE mice in terms of cell proliferation by administering BrdU 3 times (3h-apart) at 3 days post-KA injection (3dpKA) (Figure 3C). We found that the number of BrdU+ React-NSCs in the MTLE mice treated with gefitinib was significantly reduced compared to the DMSO-treated MTLE mice (drop to 47.55±9.62 %; Student’s t-test p=0.005; Supplementary Figure 3A and 3C). Although diminished BrdU+ ANPs and total BrdU cells were observed, they did not reach significance, suggesting a more specific effect of EGFR blocking in NSCs in terms of proliferation.

### HB-EGF, a natural ligand of EGFR increases early after MTLE-HS in the hippocampus

We next sought to investigate the potential natural contributors to the seizure-induced activation of EGFR in the neurogenic niche. We started exploring the expression of heparin-binding epidermal growth factor (HB-EGF), a natural ligand of EGFR that has been recently reported to be a strong mitogenic factor for astrocytes (Jia et al., 2018) and is also able to stimulate astroglial migration (Faber-Elman et al., 1996). Further, HB-EGF expression has been shown to be upregulated in the hippocampus early after KA intraperitoneal administration (Opanashuk et al., 1999). We detected increased immunolabeling for HB-EGF 3dpKA in the SGZ and the portion of the molecular layer closer to the GCL of Nestin-GFP mice (Supplementary Figure 3D). As immunohistochemical detection is not optimal for HB-EGF detection we determined its levels by ELISA at different time points (1.5h; 12h; 24h; 72h) after KA. HB-EGF showed a significant increase in the hippocampus 12hpKA and 24hpKA, decreasing to control levels at 72hpKA (876.35±33.88and 958.84±26.45 pg/mL of HB-EGF at 12h and 24h compared to 765.89±20.06 pg/mL of animals injected with saline; One-way ANOVA p=0.044 and p<0.001; Supplemental Figure 3E). In addition, cultured NSPCs costained for HB-EGF (Supplementary Figure 3F) and were able to release HB-EGF to the culture media (2152.90±202.07 pg/mL compared to 364.40 2.77 pg/mL detected in control media (without cells) and 611.40±1.39 pg/mL in the cellular pellet; Kruskal-Wallis p=0.02 and p=0.05, Supplemental Figure 3G).

### Zinc, an activator of EGFR, increases early after seizures and regulates NSPC activity *in vitro*

We next sought to investigate the potential contribution of zinc release to the activation of EGFR pathway. Zinc triggers EGFR signaling by binding directly to the receptor (Samet et al., 2003) but also indirectly by increasing HB-EGF release through the activation of zinc-dependent metalloproteinases (Wu et al., 2004). In addition, the over-release of zinc is a typical effect of neuronal hyperexcitation (Mody and Miller, 1985; Kasarskis et al., 1987). Reactive (free) zinc levels can be measured histochemically in control and MTLE-HS Nestin-GFP mice by Danscher staining (Figure 4A). Saline-injected animals showed the characteristic distribution of zinc restricted to the hilar region with an average of 107.90±9.19 zinc granules per Nestin-GFP+ cell while KA-injected animals showed a significant increase of 188.00±17.88 zinc granules per Nestin-GFP+ cell (Student’s t-test p<0.001; Figure 4B). No difference was found in the size of the granules, a measurement used as an internal control for the technique (Figure 4B). Further, because zinc homeostasis in the CNS is controlled by metallothioneins (MTH), a family of metalloproteins responsible for buffering the level of intracellular labile zinc (Kagi and Schaffer, 1988), we tested their hippocampal expression in MTLE-HS. RT-qPCR analyses determined an increase of metallothionein I (MTH1) and a tendency to increase in II (MTH2) 24-72h after KA administration (8.34±2.07 and 10.89±6.14 folds for MTH1 and 15.25±4.20 and 19.42±9.23 folds for MTH2; Kruskal-Wallis p=0.045 for MTH1 and One-way ANOVA p=0.087 for MTH2; Supplementary Figure 4A). Moreover, the presence of the proteins was confirmed by WB (Supplementary Figure 4B). No changes were detected regarding expression of MTH3. These data further support the increased presence and potential early involvement of zinc in the neurogenic niche in MTLE-HS.

**FIGURE 4.**
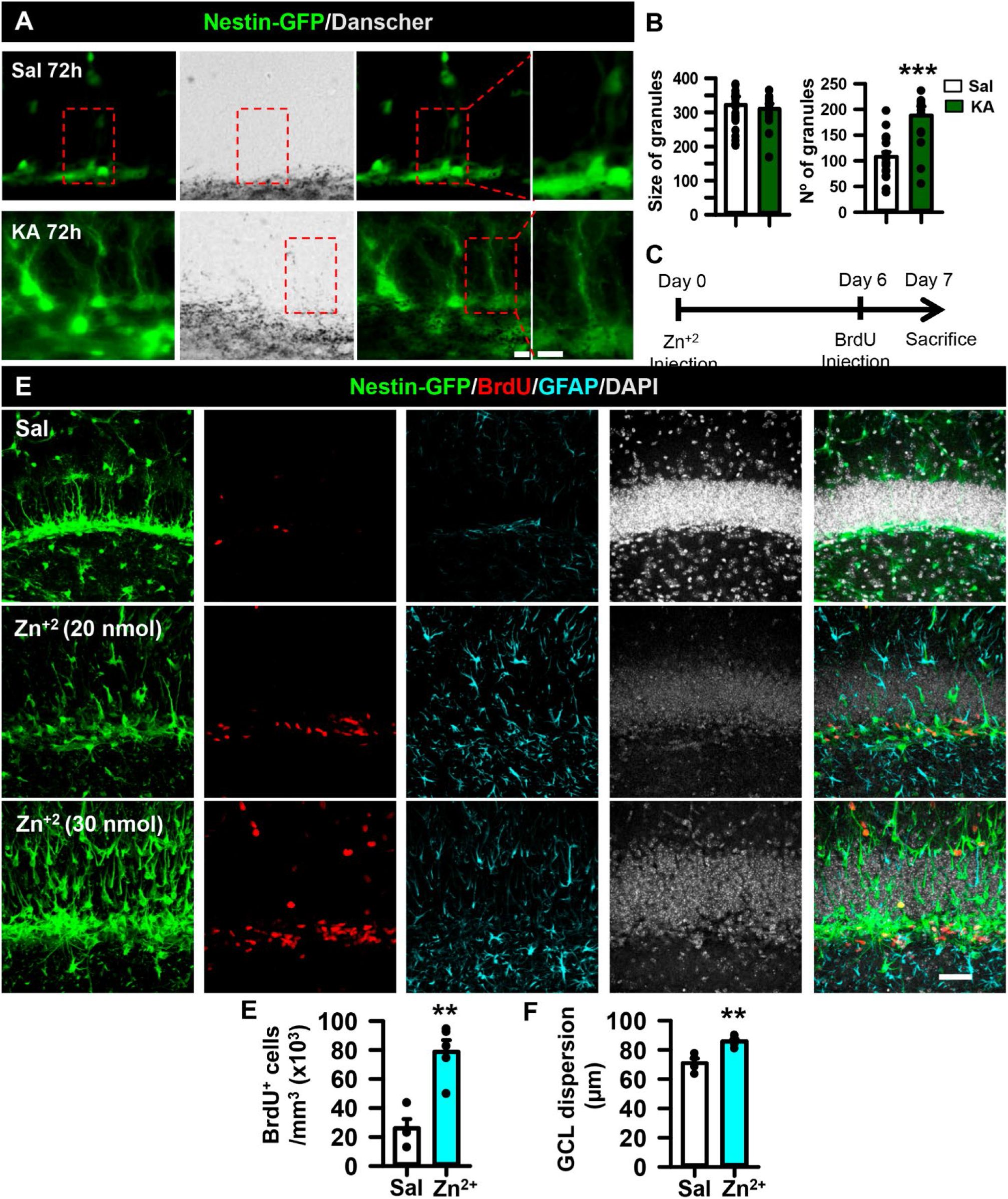
Zinc mimics the effect of seizures in the hippocampal neurogenic niche. A, Danscher staining 3 days after sal of KA intrahippocampal injection in Nestin-GFP mice showing the wider presence of zinc in the SGZ and GCL. Scale bar 10 μm. B, Quantification of the number of precipitated zinc granules. The size of the granules was assessed as an internal control. Student’s t-test ***p < 0.001. Bars show mean ± SEM. Dots show individual data. C, Schematic of the *in vivo* zinc intrahippocampal injection and BrdU administration. Nestin-GFP mice were sacrificed 7 days after zinc administration (20nmol) and BrdU was given 24h prior to sacrifice to identify mitotic cells. D, Confocal microscopy images after immunostaining for GFP, BrdU and GFAP. DAPI was used as nuclear staining. Increased cell proliferation, GCD and induction of React-NSCs are noticeable. Scale bar 10 μm. E, Quantification of the density of BrdU+ cells in the SGZ+GCL. (F) Quantification of GCD. Student’s test **p < 0.01. Bars show mean ± SEM. Dots show individual data (E, F) n=3-4.

Next, to determine the potential effects of zinc on NSPCs without the interference of the abundancy of external signals that regulate the neurogenic niche *in vivo*, we cultured NSPCs from the hippocampus of Nestin-GFP mice into laminin coated coverslips adding 0, 5, 10 and 400 μM zinc. A pulse of BrdU was given for 1h before fixation to determine cell division (Supplementary Figure 4C and 4D). We found that 5μM zinc increased cell proliferation (5512±732 BrdU+ cells compared to 2239±625 in controls; Kruskal-Wallis p=0.044; Supplementary Figure 4D). However, higher concentrations of zinc resulted in reduced cell division, an effect seemingly linked to a deleterious effect as a clear loss of cytoplasmic Nestin-GFP, as well as the abundant presence of bright and small pycnotic nuclei, was observed (Supplementary Figure 4C).

### Zinc mimics the early effect of KA on the hippocampal neurogenic niche

We then moved to an *in vivo* experiment to further corroborate the effects of zinc on the neurogenic niche. We injected intrahippocampally a single dose of 5, 20 or 30 nmol of zinc, with the first two doses reported to be tonic levels of zinc (Frederickson et al., 2006), and checked the effects after one week. 24h before sacrifice we injected BrdU intraperitoneally to label proliferating cells (Figure 4C). We did not find an effect at 5 nmol (data not shown) but GCD and more BrdU+ cells, as well as increased Nestin-GFP expression, were observed in the DG using the 20 or 30 nmol doses (Figure 4D), recapitulating some of the effects caused by the intrahippocampal injection of KA in the MTLE-HS model. We focused on the 20 nmol dose and determined first and increase in the density of BrdU+ cells that incorporated in the SGZ+ GCL, raising from 26038±6395 cells per mm^3^ to 78731±9036 cells per mm^3^ (Student’s t-test p=0.002; Figure 4E). A clear induction of React-NSCs was observed and a significant induction of GCD was measured (70.96±3.14 μm to 85.75±1.78 μm (Student’s t-test p=0.003; Figure 4F). These results show that an intrahippocampal zinc injection mimics the disruption of the neurogenic niche triggered by KA administration. As mentioned before an excess of zinc might have a positive effect by reducing neuronal hyperexcitation (Minami et al., 2006) and therefore in our context of interest it could be argued that zinc chelation could be used to preserve the neurogenic niche in MTLE-HS. To test this hypothesis, we administered subcutaneously 5 mg/Kg of the zinc chelating agent TPEN as previously described (Kim et al., 2012) on both control and MTLE-HS Nestin-GFP mice twice a day during 7 consecutive days starting the same day that saline or KA was injected into the hippocampus to induce MTLE-HS. Although we did not find evident variations on GCD irrespective of the treatment (Supplementary Figure 4E) we quantified the number of dying cells by condensed DAPI staining, expecting a significant increase after KA administration as previously reported (Sierra et al., 2015b) and evaluating the effect of zinc chelation by TPEN (Supplementary Figure 4F). We observed a tendency in the GCD and cell death to increase after KA compared to saline-injected animals in both vehicle and TPEN treated mice. However, TPEN failed to reduce cell death, as a similar density of pycnotic nuclei was found in the GCL than KA+vehicle animals and significantly higher than saline+TPEN animals (24864.24±630.84 pycnotic nuclei per mm^3^ for KA+TPEN respect to 2895.61±1635.08 pycnotic nuclei per mm^3^ for Saline+TPEN; Kruskal-Wallis p=0.014; Supplementary Figure 4F). These results suggest that despite mimicking the early effects of MTLE-HS on the neurogenic niche, endogenous zinc also exerts a neuroprotective effect.

### HB-EGF and zinc activate the EGFR pathway in NSPCs *in vitro*

In order to confirm that the presence of zinc could modulate NSPCs in an EGFR-mediated manner, we again cultured NSPCs from the hippocampus of Nestin-GFP mice. To avoid EGFR activation caused by the growth factors present in the culture media, NSPCs were subjected to growth factor starvation during 15 min, 2h or 6h. A marked loss of phosphorylated-ERK (p-ERK) was found at the 2h time point (Supplementary Figure 5A). Following an adaptation of a protocol previously described (Wu et al., 2004), NSPCs were exposed to either 100 ng/mL EGF, 200-μM zinc, or pretreated with 2-μM gefitinib for 60 min prior to the addition of 200-μM zinc. Protein extracts were blotted sequentially against Tyr845 and Tyr1068 phosphorylation sites of EGFR and to total EGFR to test receptor phosphorylation (Figure 5A and Supplementary Figure 5B). Although we could not fully exclude that autocrine-released HB-EGF could trigger some residual EGFR phosphorylation, the starving condition was considered the control as with the lowest phosphorylation (ratio to control=1). We observed that zinc was able to induce a mild phosphorylation of Tyr845 that was fully blocked in the presence of gefitinib (ratio 1.00±0.13 starved compared to 2.36±0.20 zinc; Kruskal Wallis p=0.05; and 2.36±0.20 zinc compared to 0.42±0.04 zinc+gefitinib; Kruskal-Wallis p=0.02; Figure 5B and 5C). There was not, however, statistical difference for Tyr1068 phosphorylation (Supplementary Figure 5C). It has also been reported that an opposite feed-back effect by which p-ERK is able to mediate HB-EGF shedding and subsequent EGFR activation can take place (Yin and Yu, 2009). We observed that zinc exposure in starved conditions was able to increase p-ERK signaling and interestingly, the inhibition of EGFR reduced its phosphorylation (ratio 1.00±0.04 starved compared to 2.23±0.16 zinc; One-way ANOVA p=0.017; and 2.23±0.16 zinc compared to 0.85±0.35 zinc+gefitinib; One-way ANOVA p=0.015; Figure 5B and 5D). To further corroborate this effect and to get closer to *in vivo* conditions, we stimulated starved cells with the highly mitotic HB-EGF ligand, HB-EGF+zinc, or we pretreated cells with gefitinib and then exposed to HB-EGF+zinc. Our results demonstrated that HB-EGF was able to strongly phosphorylate EGFR at both Tyr845 and Tyr1068 residues. Remarkably, zinc stimulation did not produce any significant additive effect, possibly due to the excess of HB-EGF. Importantly, gefitinib pretreatment was able to block phosphorylation at Tyr845 and Tyr1068 (ratio 1.00±0.29 starved compared to 3.12±0.52 zinc+HB-EGF; One-way ANOVA p=0.027; and 3.12±0.52 zinc+HB-EGF compared to 1.09±0.39; zinc+HB-EGF+gefitinib; One-way ANOVA p=0.046 for Tyr845; and ratio 1.00±0.16 starved compared to 4.63±0.89 zinc+HB-EGF; One-way ANOVA p=0.005; and 4.63±0.89 zinc+HB-EGF compared to; 0.51±0.14; zinc+HB-EGF+gefitinib; One-way ANOVA p=0.002 for Tyr1068; Figure 5E, 5F and 5G). Moreover, EGFR inhibition showed a tendency to reduce downstream p-ERK even in the concomitant presence of both zinc and HB-EGF (Figure 5E and 5H). These results suggested that inhibiting EGFR with gefitinib was sufficient to inactivate the receptor regardless of the autocrine and paracrine HB-EGF-dependent phosphorylation.

**FIGURE 5.**
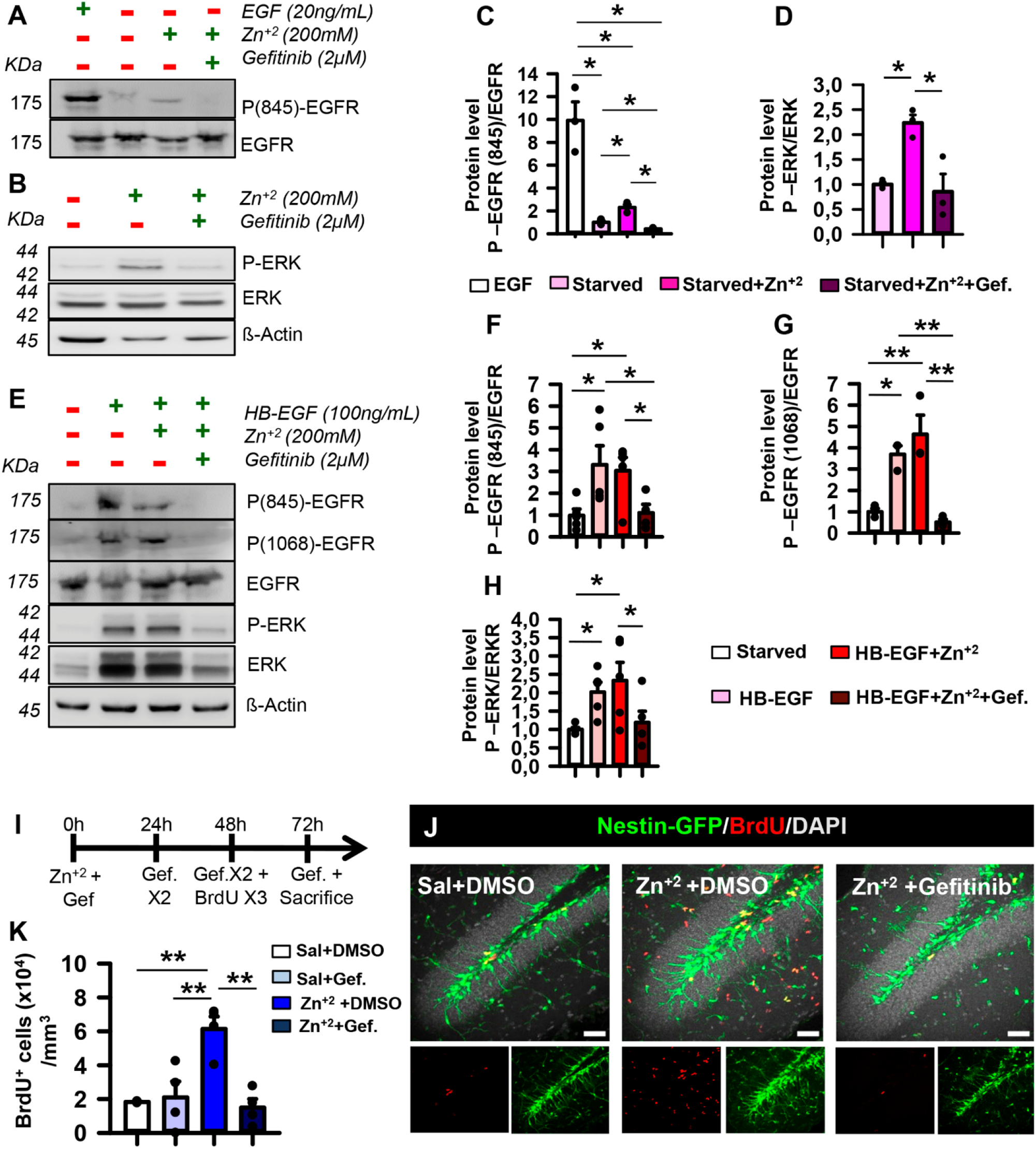
Gefitinib blocks the zinc-induced activation of the EGFR signaling pathway in NSPCs. A, Pretreatment with gefinitib (2μM) prevents the phosphorylation of EGFR induced by EGF and by zinc (200μM) as shown by WB. B, WB of *in vitro* cultured NSPCs showing that activation of p-ERK downstream signaling in presence of zinc (200μM) is blocked by gefitinib (2μM). Note that β-Actin shows no variation on loading inputs. C, Ratio of phosphorylated to non-phosphorylated EGFR Tyr845 after Kruskal-Wallis analysis. D, Ratio of phosphorylated to non-phosphorylated ERK after one-way ANOVA. *p < 0.05. Bars show mean ± SEM. Dots show individual data (C, D). E, the presence of HB-EGF and zinc stimulates EGFR phosphorylation at Tyr845 and Tyr1068 sites and shows a tendency to stimulate p-ERK signaling. Gefitinib reduces both phosphorylation of EGFR and p-ERK. F, Quantification of the phosphorylated to the non-phosphorylated ratio EGFR Tyr845. G, Quantifications of the phosphorylated to the non-phosphorylated ratio EGFR Tyr1068 followed by One-way ANOVA analysis (F, G). (H) Quantifications of the phosphorylated to the non-phosphorylated ratio for ERK followed by Kruskal-Wallis analysis. *p < 0.05, **p < 0.01. Bars show mean ± SEM. Dots show individual data (F-H). I, Schematic of intranasal gefitinib treatment after intrahippocampal zinc injection. BrdU was given 24h prior to sacrifice to identify mitotic cells. J, Confocal microscopy images of saline, zinc and zinc+gefitinib (10mg/Kg) 3d after zinc showing the increase of BrdU+ cells in the SGZ of zinc-injected animals, that is reversed by the administration of gefitinib. Scale bar 50 μm. K, Quantification of total BrdU+ cells in the SGZ. There is a significant increase of BrdU+ cells in the SGZ after zinc administration, that is however contained administering gefitinib. One-way ANOVA *p < 0.05. Bars show mean ± SEM. Dots show individual data. n=3.

### Zinc induces React-NSCs through activation of EGFR *in vivo*

The next step was to check the *in vivo* effect of EGFR inhibition in animals that received a single dose of zinc (20 nmol) intrahippocampally, expecting that blocking EGFR should reduce the zinc-induced over-proliferation of cells. Towards this purpose we administered gefitinib intranasally during the first 72h following the same paradigm described before. The animals received three doses (3h-apart) of BrdU intraperitoneally to track cells under division starting 24h before sacrifice (Figure 5I). Our results showed that indeed EGFR inhibition reduced the increase of BrdU+ cells in the GCL provoked by the intrahippocampal injection of zinc (15680±2590 saline compared to 51360±7890 zinc; One-way ANOVA p=0.016; and 51360±7890 zinc compared to 23970±6830 zinc+gefitinib; One-way ANOVA p=0.026; Figure 5J-K). No significant changes in expression and distribution of EGFR by NSCs was found 24h after zinc injection (Supplementary Figure 5D and E).

To check the effect on early neurogenesis we repeated the experiment but sacrificed the animals at 14d post-intrahippocampal zinc injection. Despite the increase in proliferation and the drastic change of the neurogenic niche in the short-term, similar to that of MTLE, we observed no ablation of DCX+ cells (Supplementary Figure 5D-E). On the contrary, the results showed a tendency of neurogenesis to increase, expressed as the ratio of BrdU+DCX+ cells to DCX+ cells (Supplementary Figure 5D and 5F). These results support the notion that zinc is another contributor to the complex responses triggered by seizures rather a single master effector. As an example, no significant changes in expression and distribution of EGFR by NSCs was found 24h after zinc injection (Supplementary Figure 5G-H).

## DISCUSSION

### Seizures induce the transformation of hippocampal NSCs into gliogenic React-NSCs

A population of radial glia-like NSCs were shown to be the source of postnatal and adult neurogenesis in the hippocampus in a pioneering work that also hinted at the possibility that they could generate astrocytes as well (Seri et al., 2001). This astrogliogenic capability was later proved (Encinas et al., 2011), together with their capacity to also generate more NSCs (Bonaguidi et al., 2011; Pilz et al., 2018). Thus, the natural multipotent capability of NSCs in natural conditions was established. More recently the cell-differentiation diversity of hippocampal NSCs was further expanded as we showed that seizures triggered the conversion of NSCs into reactive astrocytes through an intermediate cell type, different from both NSCs and reactive astrocytes, termed React-NSCs (Sierra et al., 2015b; Muro-García et al., 2019). Similarly, it was reported a few months later that NSCs from the SVZ generated reactive astrocytes that migrated into the cortex in which stroke had been experimentally induced (Faiz et al., 2015). Complete ablation of neurogenesis might be a particular outcome of MTLE with HS as in other models a boost of neurogenesis, with abnormal features (aberrant neurogenesis), is generally found (Kuruba et al., 2009; Pineda and Encinas, 2016). A straightforward comparison among different models (intracerebral KA, intraperitoneal KA, pilocarpine, electrical kindling…) cannot be performed because of the different paradigms and points of analysis of cell markers, cell proliferation, cell survival, cell differentiation and the different cell types studied (NSCs, newborn neurons, astrocytes…). The mechanisms driving increased aberrant neurogenesis in MTLE versus the absence of neurogenesis in MTLE-HS deserve further attention on their own. For the moment we can speculate that the different levels of local neuronal hyperexcitation and the presence or absence of neuronal death and reactive gliosis in the hippocampus can trigger qualitatively different responses in the neurogenic nice, which reflects the high level of plasticity of NSCs and neurogenesis (Bielefeld et al., 2019).

Characterizing the active role of NSCs in the brain’s response to damage by contributing to reactive gliosis opens new venues for potential therapeutic interventions. Unveiling the mechanisms controlling the transformation of NSCs to React-NSCs might allow us to not only preserve healthy NSCs and eventually neurogenesis in MTLE-HS but also to manipulate the level of reactive gliosis. We herein show that the EGFR is a key regulator in the induction of React-NSCs after seizures, and that its blockage can preserve neurogenesis. Further, we show that not only HB-EGF, the natural ligand of EGFR, but also zinc, is involved in the EGFR-dependent induction of React-NSCs.

### Intrahippocampal KA injection triggers the overexpression of EGFR in hippocampal NSCs. Blocking EGFR with Gefitinib partially preserves NSCs

*In vitro* studies using stem cell-like germinative cells highlighted the EGFR/ERK transduction pathway as responsible to activate and induce proliferation upon addition of human EGF with EGFR inhibition blocking cell proliferation (Cheng et al., 2017). Other works demonstrated that the inactivation of the EGFR/MAPK pathway blocked or greatly retarded NSC cycle progression in Droshophila (Li et al., 2015), and the absence of EGFR strongly impaired stem cell self-renewal (Robson et al., 2018). EGFR has been shown to be expressed in activated and proliferating NSPCs (Okano et al., 1996; Jhaveri et al., 2015) and it is actually used to separate the activated NSPCs from quiescent ones (Walker et al., 2016). In addition, the activation of the EGFR controls the transformation of astrocytes into hypertrophic reactive astrocytes (Liu and Neufeld, 2004; Liu et al., 2006; Tsugane et al., 2007) acting also as an upstream signal to exit quiescence (Liu et al., 2006). Thus, we thought that EGFR was a good candidate to be regulating the transformation of NSCs into React-NSCs, a transitory cell-type different from both NSCs and reactive astrocytes but with common features to both cell types. Thus, we first confirmed that early after the induction of MTLE-HS, EGFR was expressed in higher levels in the hippocampus and in particular in the neurogenic niche, in which it colocalized with Nestin-GFP+ NSCs and ANPs. Downstream effectors of the EGFR, such as phospho-STAT3 and phospho-ERK1/2, were also elevated. Blocking the EGFR in hippocampus-derived NSPCs with gefitinib reduced cell proliferation, a result that was replicated *in vivo*. Importantly, gefitinib not only reduced the seizure-induced increase in NSCs proliferation but also, at least partially, preserved the healthy morphology of NSCs.

### HB-EGF, a natural ligand of EGFR, is also overexpressed after seizures and its interplay with over-released zinc contribute to EGFR activation after seizures

We studied the role of HB-EGF, a natural ligand of EGFR. Binding of HB-EGF to EGFR has been associated with enhanced EGFR tyrosine kinase activity and prolonged ERK activation (Yoo et al., 2012). Our results also show that an early increase in the expression of HB-EGF, the natural ligand of EGFR, takes place in parallel to the increase of EGFR expression. This result is in agreement with previous work based on intraperitoneal administration of KA, in which HB-EGF mRNA increased within 3h and continued elevated at least 48h in the DG (Opanashuk et al., 1999). EGFR and HB-EGF have been associated with pro-inflammatory cytokine release (Richter et al., 2002) and HB-EGF induced astrocyte proliferation *in vitro* in an EGFR dependent manner. Thus, HB-EGF could be also a potential therapeutic target but because it also enhances neuronal and astrocyte survival (Kornblum et al., 1999; Jia et al., 2018) and is neuroprotective against excitotoxicity (Opanashuk et al., 1999), this option is not encouraging.

We were intrigued by the potential role of zinc in the activation of the EGFR pathway for several reasons. 1) HB-EGF is initially synthesized as a membrane-bound precursor (pro-HB-EGF), it is cleaved at the juxtamembrane domain to release the soluble form of HB-EGF by matrix metalloproteinases (Izumi et al., 1998), and these are activated by zinc (Le Gall et al., 2003; Wu et al., 2004). 2) Zinc-dependent ERK activation can stimulate EGFR phosphorylation through HB-EGF (Yin and Yu, 2009). 3) Zinc is massively released from excitatory terminals during neuronal hyperactivity reducing neuronal activation (Assaf and Chung, 1984). In the brain the most abundant concentration of zinc has been found in the Mossy fibers (MFs) of granule cells(Frederickson, 1989). MFs connects to CA3 pyramidal neurons and dentate hilar cells. However, after KA lesion, the MFs sprout in the DG forming new synapses on granule cell dendrites, which increase the excitatory connections between granule cells (Buckmaster et al., 2002) and seizure susceptibility(Sutula et al., 1989; Shetty et al., 2005). Knockout mice lacking the synaptic zinc transporter have increased excitotoxicity after KA-induced neuronal hyperexcitation (Cole et al., 2000). Additionally, zinc chelation was able to provoke the induction of seizures producing and paroxysmal epileptiform brain activity and subsequent damage(Mitchell and Barnes, 1993; Cuajungco and Lees, 1998). Further, the chelation of zinc during neural hyperexcitation exacerbated excitotoxicity (Domínguez et al., 2003, 2006). Other works suggested that zinc release into extracellular space by glutamatergic synapses could be toxic (Babb et al., 1991; Lee et al., 2000). Our data reported herein support the detrimental effect of zinc chelation, as it exacerbated cell death in MTLE-HS. Nevertheless, we found that intrahippocampal zinc administration recapitulated several of the effects of the KA injection as a model of MTLE-HS. The intrahippocampal injection of zinc triggered massive proliferation and reactive gliosis, including the induction of React-NSCs in the neurogenic niche and GCD in a dose dependent manner. Interestingly, the extracellular matrix has an important role regulating physiological plasticity during epileptogenesis (Dityatev and Fellin, 2008). Extracellular matrix disintegrins and metalloproteases contain zinc-binding motif critical for proteinase activity(Sagane et al., 1998). It is plausible to speculate that an alteration of its function could affect the proteolytic cleavage of proteins such as Reelin, that has a direct role in neuronal migration affecting thus to GCD (Tinnes et al., 2011). In cultured NSPCs zinc (and HB-EGF) increased cell proliferation and activated EGFR and its transduction pathway. To test the hypothesis that zinc could be directly acting on EGFR we used again gefitinib *in vitro* and *in vivo*. As gefitinib indeed blocked the effects of zinc on cultured NSPCs *in vitro* and in the neurogenic niche *in vivo*, we can conclude that the excess of zinc released during seizures contributes to the induction of React-NSCs by directly activating the EGFR that NSCs overexpress due to neuronal hyperexcitation.

Regulating zinc concentration is therefore a key event that can dictate the final effect of zinc in neuronal hyperexcitation. Intracellular zinc buffering is largely regulated by a family of small cysteine-rich proteins called metallothioneins (MTH) (Ebadi and Hama, 1986). We found that MTH-1 mRNA was significantly increased at 24h post KA administration and a tendency to increase was observable for MTH-2 from 12h onwards, highlighting that the triggering of environmental changes mediated by the zinc release occurs very early on time, and probably play a role for the shedding and release of diffusible HB-EGF.

### Blocking EGFR directly, rather than targeting HB-EGF or zinc, is a better strategy for preserving NSCs

Overall, our and former results strongly suggest that targeting directly EGFR rather than the release of zinc or HB-EGF is a better strategy for potential therapeutic benefit of preserving NSCs and neurogenesis. Treatment with EGFR inhibitors has been reported as neuroprotective in rat models of glaucoma (Liu et al., 2006) and spinal cord injury (Erschbamer et al., 2007) by acting preferentially on reactive astrocytes. Our results showed that EGFR inhibition was able to block ERK1/2 phosphorylation on NSPCs, suggesting the participation of EGFR in the mitogenic ERK1/2 pathway on NSC. Although gefitinib strongly impaired cell division assessed by BrdU incorporation on treated NSPC cultures, we still observed a fraction of cells undergoing active nucleotide incorporation and we cannot discard the concomitant involvement of other signaling pathways. Ablation of cell division by irradiation in the SVZ enriched the fraction of EGFR-negative quiescent NSCs. However, NSCs were able to enter into the cell cycle shortly after (24-48h) to repopulate the niche (Daynac et al., 2013) suggesting that other signaling pathways such as Sonic hedgehog (Shh) (Daynac et al., 2016) also play an important role in NSC activation. In conclusion, in the present work we show for the first time a molecular mechanism involved in the induction of React-NSCs and the subsequent loss of neurogenesis: the EGFR pathway. After seizures EGFR expression increases specifically in NSCs in the hippocampal neurogenic niche, as does that of HB-EGF and the release of zinc. HB-EGF is a natural ligand of the EGFR and zinc can bind directly to the EGFR to activate it. But zinc can also potentiate the release of HB-EGF, thus further contributing to EGFR activation. By blocking EGFR with gefitinib, we were able to block the induction of React-NSCs, at least in terms of activation and morphology.

## ACKNOWLEDGEMENTS

We thank the staff at the Leioa animal facility of UPV/EHU, Laura Escobar at the Imaging Core Facility in Achucarro and the whole Laboratory of Neural Stem Cells and Neurogenesis Laboratory for insight and discussion. This work has been funded by Spanish Ministry of Economy and Competitiveness (MINECO, with FEDER Funds) grants SAF2-015-70866-R and PID2019-104766RB-C21 to J.M.E. and J.R.P,; by the MINECO Ramón y Cajal Program: RYC-2013-13450 to J.R.P. and RYC 2012-11137 to J.M.E.; and by the MINECO PCIN-2016-128 (ERA-NET-NEURON III program) to J.M.E.; O.P.-A. holds a UPV/EHU predoctoral fellowship, I.D. holds a FPI (MINECO) predoctoral grant and S.M.-S. held a Fundación Gangoiti predoctoral fellowship.

## AUTHOR CONTRIBUTIONS

Conceptualization, O.P.-A., J.M.E., and J.R.P.; Methodology, O.P.-A., I.D., S.B.-C., E.V., S.M.-S., and J.R.P.; Formal Analysis, O.P.-A., I.D., S.B.-C., E.V., S.M.-S., and J.R.P.; Investigation, O.P.-A., J.M.E., and J.R.P.; Data Curation, O.P.-A., I.D., S.B.-C., E.V., S.M.-S., J.M.E., and J.R.P.; Writing – Original Draft, O.P.-A., and J.R.P. Writing – Review and Editing, O.P.-A., J.M.E., and J.R.P.; Visualization, O.P.-A., J.M.E., and J.R.P.; Funding acquisition, J.M.E., and J.R.P.; Supervision, J.M.E., and J.R.P.

## DATA AVAILABILITY STATEMENT

The data that support the findings of this study are available from the corresponding authors upon reasonable request.

## ETHICAL PUBLICATION STATEMENT

We confirm that we have read the Journal’s position on issues involved in ethical publication and confirm that this report is consistent with those guidelines.

## SUPPLEMENTARY FIGURES

**Figure S1.**
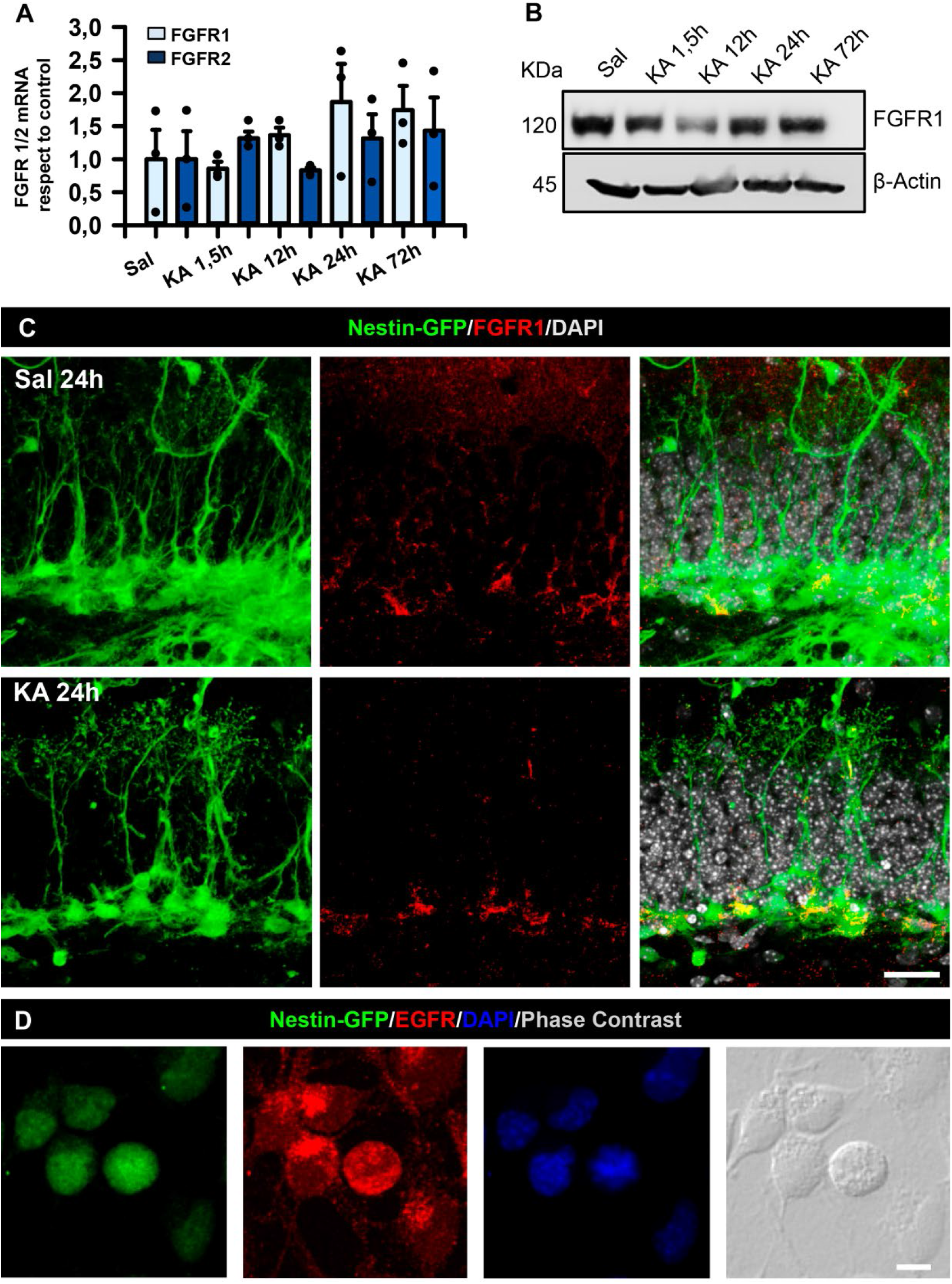
FGFR is not increased in the DG of MTLE mouse model. (A) RT-qPCR determination of changes of FGFR1 and FGFR2 mRNA expression during the initial 3d after intrahippocampal injection of KA (1 nmol). One-way ANOVA for FGFR1 and Kruskal Wallis for FGFR2. Bars show mean ± SEM. Dots show individual data. n=3. (B) WB of hippocampi dissected at different time points after KA (1 nmol) showing no changes in FGFR1 expression. (C) Confocal microscopy images after immunostaining for FGFR1 in the SGZ of Nestin-GFP mice 24 h after KA. Scale bar is 10 μm. (D) Immunofluorescence images of cultured hippocampal NSPCs stimulated with EGF and FGF showing EGFR staining. Scale bar is 10 μm.

**Figure S2.**
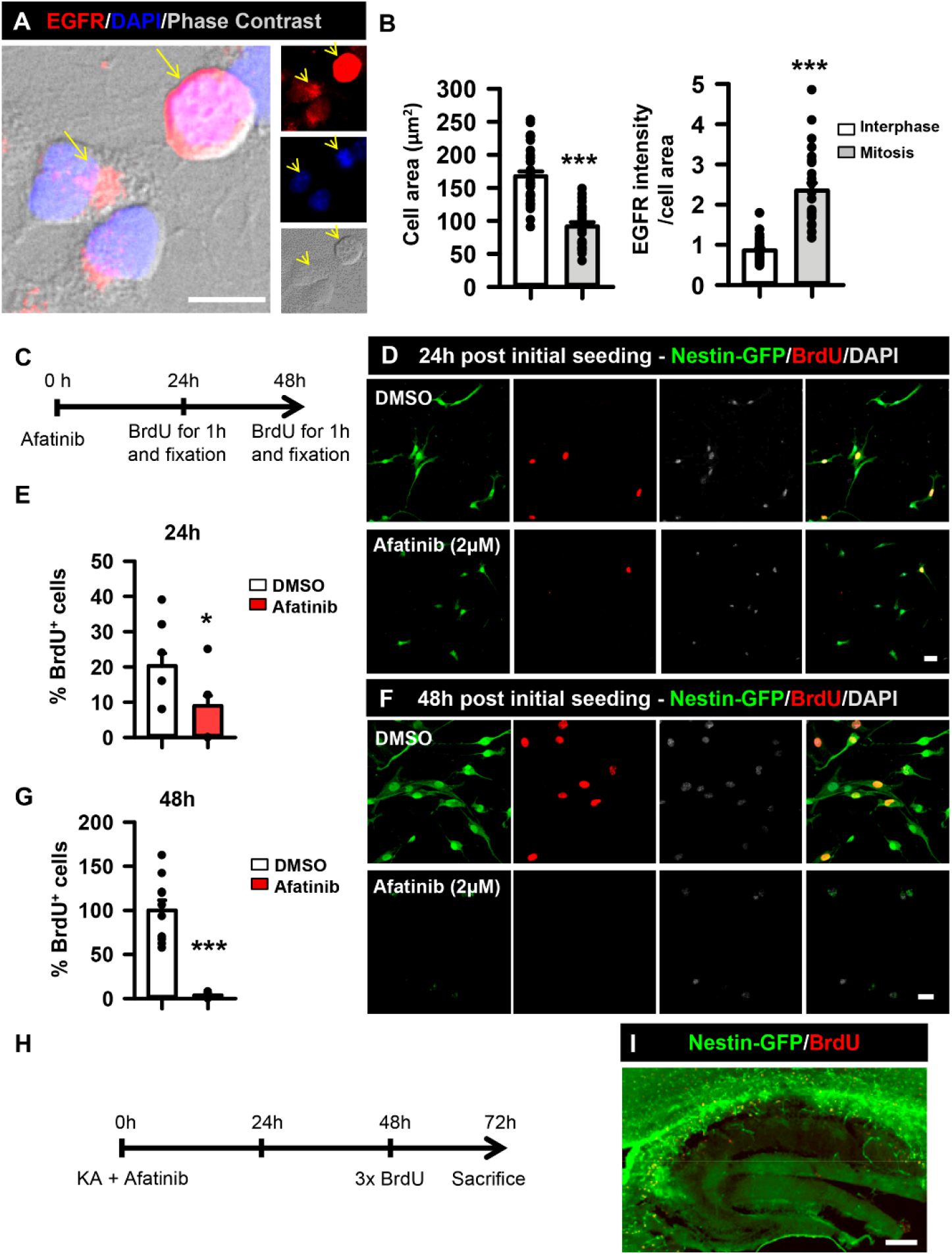
EGFR increases in mitotic cells and its irreversible inhibition provokes cell death both *in vitro* and *in vivo*. (A) Immunofluorescence images showing the increase in EGFR expression in cultured hippocampal NSPCs during mitosis. Arrows mark cell localization. Scale bar 10 μm. (B) Quantification of the cell area and pixel intensity (EGFR staining) of cultured NSPCs undergoing mitosis or in interphase. Student’s t-test for cell area and Mann Whitney for EGFR pixel intensity ***p < 0.001. Bars show mean ± SEM. Dots show individual data. (C) Schematic of the culture strategy to evaluate the effect of the irreversible EGFR inhibitor afatinib on cell proliferation. NSPCs were cultured in presence of EGF or afatinib pretreatment plus EGF during 48h. A pulse of BrdU 10 μM 1 h before fixation was applied to label mitotic cells. (D) Representative immunofluorescence images of cultured NSPCs from Nestin-GFP mice showing the reduction of nuclear BrdU incorporation 24 h post afatinib treatment. Scale bar is 20 μm. (E) Quantification of BrdU+ cells expressed as percentage of total number of cells. Mann Whitney *p < 0.05. Bars show mean ± SEM. Dots show individual data. (F) Representative immunofluorescence images of cultured NSPCs 48 h post-afatinib treatment, illustrating the cell loss. Scale bar is 20 μm. (G) Quantification of BrdU+ cells showing the drastic reduction due to afatinib treatment. Mann Whitney ***p < 0.001. Bars show mean ± SEM. Dots show individual data. (H) Schematic of the *in vivo* strategy to assess the effect of afatinib on the neurogenic niche. Afatinib or vehicle was administered together with KA and mice were sacrificed 3 d later. (I) Confocal microscopy image showing the massive cell death provoked in the hippocampus by afatinib administration in a Nestin-GFP mouse, supporting the *in vitro* results.

**Figure S3.**
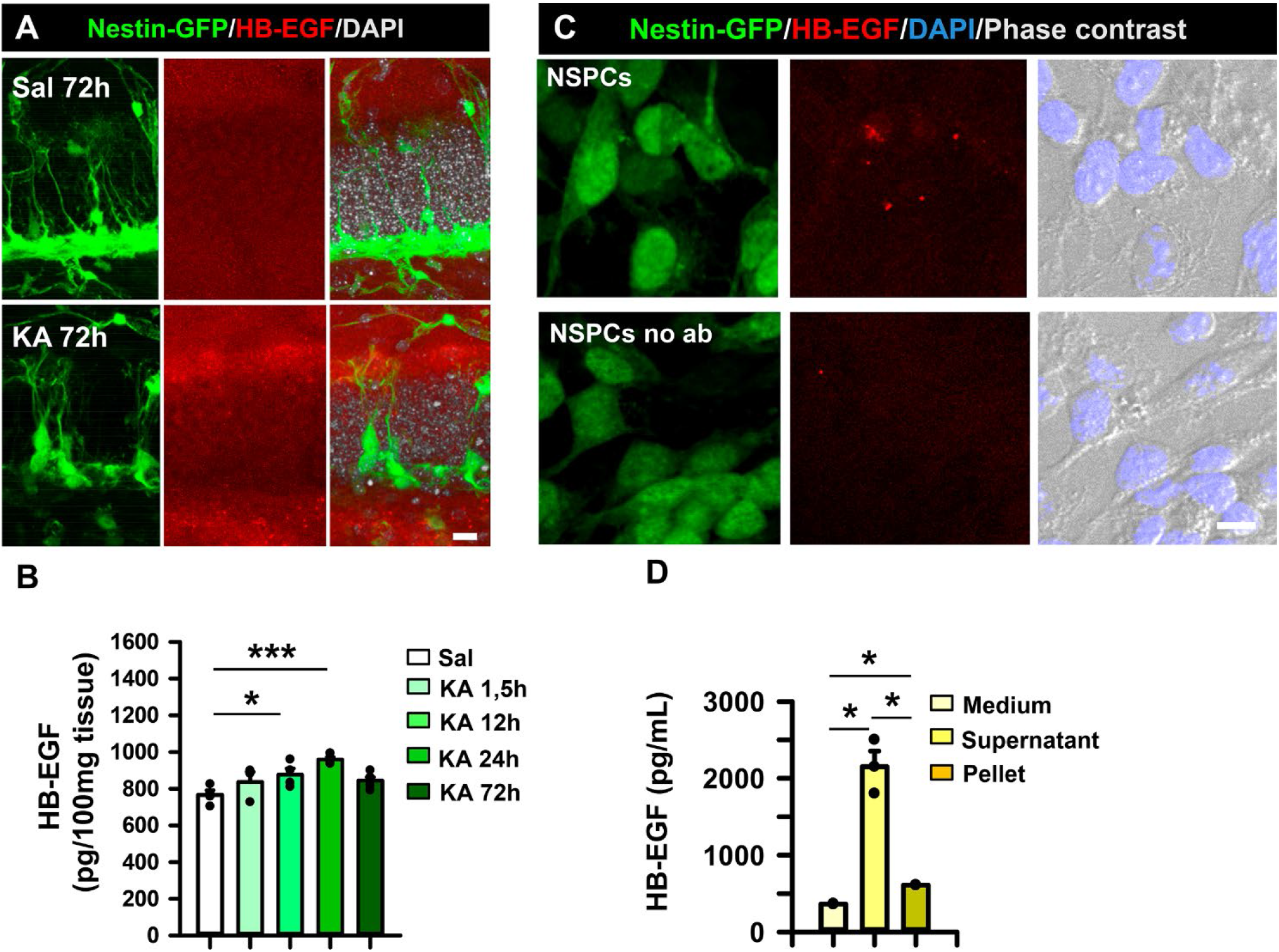
Pharmacological blockage of EGFR signaling 14dpKA. HB-EGF increases early after the induction of MTLE *in vivo* and is secreted by NSPCs *in vitro*. (A) Confocal microscopy images showing an increase of the EGFR ligand HB-EGF in the hilus and the molecular layer of Nestin-GFP mice 3dpKA. Scale bar 20 μm. (B) Quantification of HB-EGF by ELISA at different time points during the first 3dpKA. Values are represented as pg of HB-EGF in 100 μg of tissue. One-way ANOVA *p < 0.05, ***p < 0.001. Bars show mean ± SEM. n=3. Dots show individual data. (C) Immunofluorescence images of NSPCs *in vitro* showing HB-EGF expression. Scale bar is 10 μm. (D) Quantification by ELISA of HB-EGF released *in vitro*,expressed as pg of ligand per mL. Kruskal Wallis *p < 0.05. Bars show mean ± SEM. Dots show individual data.

**Figure S4.**
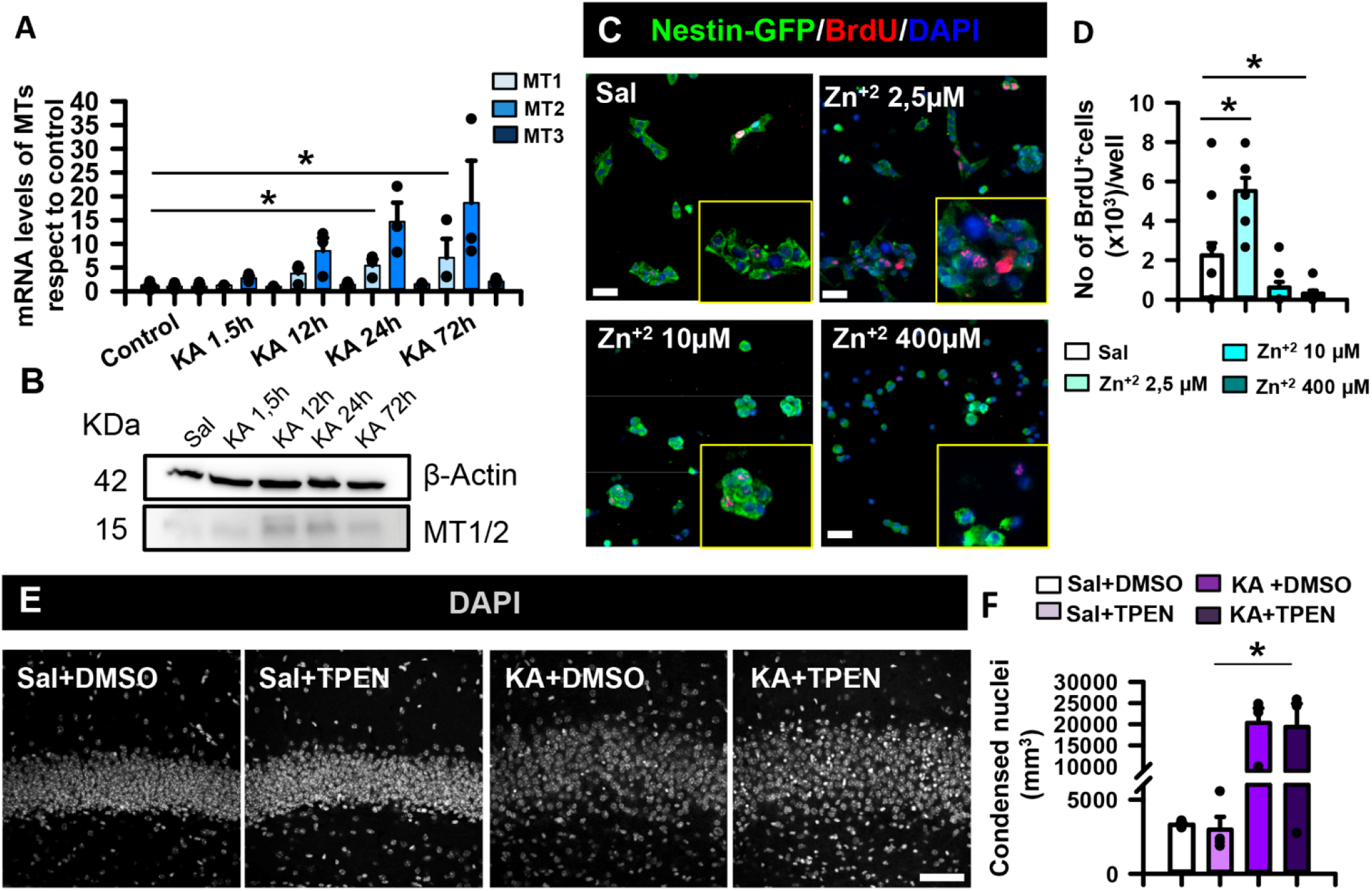
Zinc recapitulates niche disorganization and increase the number of proliferating cells. However, its chelation increases cell death. (A) RT-qPCR determination of changes in MTH I, II & III mRNA expression during the initial 3d after KA (1nmol) intrahippocampal injection. Kruskal-Wallis for MT1 and One-way ANOVA for MTH2 and MTH3 *p < 0.05. Bars show mean ± SEM. Dots show individual data. (B) WB of MTH I and II proteins at different time points after KA injection. (C) Immunofluorescence images of NSPCs in the presence of 5, 10 and 400 μM of zinc for 24h. Note that 400 μM induces cell death. A pulse of BrdU 10 μM was given 1h before fixation to label mitotic cells. Scale bar 20 μm. (D) Quantification of the number of BrdU+cells in the cultured NSPCs treated with zinc. Kruskal-Wallis *p < 0.05. Bars show mean ± SEM. n=3. Dots show individual data. (E) Confocal microscopy images showing the effect of the use of TPEN, a zinc chelator, in saline and KA-injected animals. Zinc chelation exacerbated nuclei dispersion and accumulation of condensed pycnotic nuclei as seen after DAPI staining. (F) Quantification of condensed pycnotic nuclei per mm^2^ in the DG after chelation of zinc. Kruskal-Wallis *p < 0.05. Bars show mean ± SEM. Dots show individual data.

**Figure S5.**
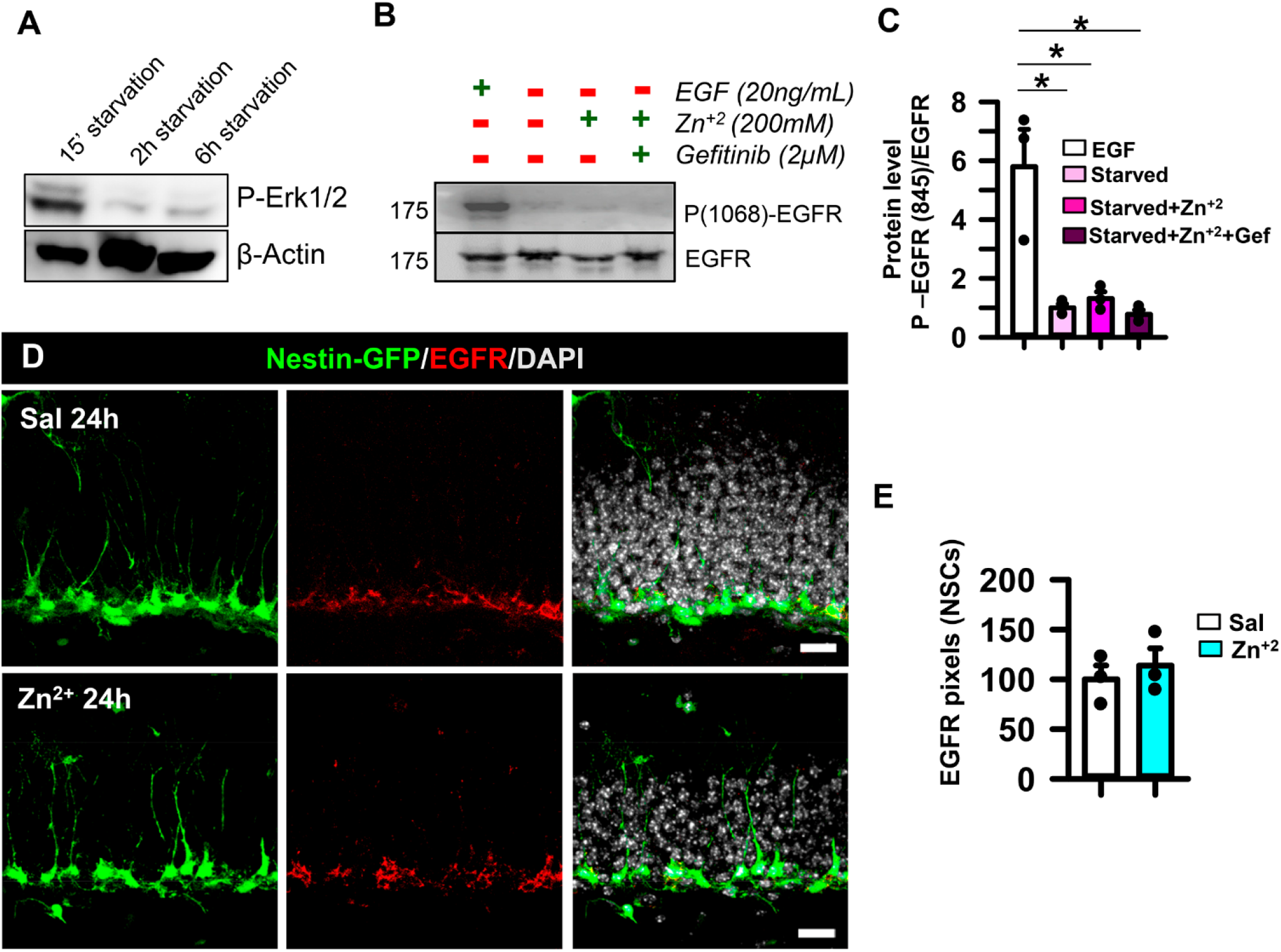
Zinc does not induce the phosphorylation of EGFR Tyr1068, does not alter EGFR expression. (A) WB for phosphorylated-ERK of cultured NSPCs after 15 min, 2 h and 6 h of growth factors starvation. (B) WB for EGFR and phosphorylated-1068-EGFR of cultured NSPCs stimulated either with EGF (20ng/mL) or starved and stimulated with zinc (200μM), or zinc with gefitinib (2μM) pretreatment. (C) Quantification of the ratio of phosphorylated-EGFR to EGFR. Kruskal-Wallis, *p < 0.05. (D) Confocal microscopy images of the neurogenic niche from Nestin-GFP mice after immunostaining for EGFR 24h post intrahippocampal zinc injection. Scale bar 20 μm. (E) Quantification of EGFR expression (in pixels) in NSCs represented as the percentage respect to control. Student’s t-test. Bars show mean ± SEM. Dots show individual data. n=3.

## 1. MATERIALS and METHODS

### 1.1 Mice

Experimental procedures were performed in compliance with the European Community Council Directive of November 24, 1986 (86/609/EEC) and were approved by the University of the Basque Country (UPV/EHU) Ethics Committees (Leioa, Spain) and the Diputación Foral de Bizkaia. The protocol reference is CEEA M20/2015/236. The animals were maintained with access to food and water ad libitum in a colony room that was maintained at a constant temperature (19–22°C) and humidity (40–50%) on a 12:12 h light/dark cycle. Nestin-green fluorescent protein (GFP) (Mignone et al., 2004) and wild type (WT) Nestin-GFP-negative littermates) on a C57BL/6j background counterparts were used. The animals were 2 months old and weighted 25-30 g at the time of experimentation. Both male and females were indistinctly used in each experiment and littermates were randomly assigned to each experimental condition.

### 1.2 Cell Cultures

Adult hippocampal neural stem and progenitor cell (NSPC) cultures were obtained using an adaptation of protocols previously described (Pineda et al., 2013; Jhaveri et al., 2015). NSPCs were isolated from either Nestin-GFP or WT littermates. Briefly, hippocampi were dissected from 5 adult mice and placed in ice-cooled PBS with sucrose (PBS: Phosphate-buffered saline; Cat#806544 & sucrose; Cat# S0389, Sigma-Aldrich, St Louis, MO, USA). The tissue was cut in chunks of 2-3mm size that were pooled together and incubated with a mixture of papain (1 mg/ml papain (15 UI/ml), Cat# LS003126; Worthington, Lakewood, NJ, USA) DNAse (2.5μl DNAse 2000 U/mL, Cat# 10636153 Fisher Scientific, Waltham, Massachusetts, USA) for 30 min for enzymatic digestion. Enzymes were inactivated with Ovomucoid solution (0.7 mg/ml, Cat#T9253, Sigma-Aldrich, St Louis, MO, USA) and tissue was mechanically homogenized with a P1000 pipet tip. The resultant cell mixture was passed through a 20 μm Nylon filter (Filcons, Cat# Z290823, Agilent Technologies, Santa Clara, CA 95051, USA) and washed twice with bovine serum albumin (BSA) diluted with PBS at 0.15%. The cell mixture underwent centrifugation at 200g for 10min and then cultured using NeuroCult proliferation medium (Cat#05702, Stem Cell Technologies, Vancouver, Canada). The cells were maintained at standard conditions in a humidified 37 C° incubator containing 5% CO2. NSPC cultures were then passaged every 7 days by enzymatic disaggregation with Accutase (cat# 7920, Stem Cell Technologies, Vancouver, Canada). The medium was supplemented with Neurocult proliferation supplement (cat# 05702, Stem Cell Technologies, Vancouver, Canada) at 9:1 ratio; Heparin solution 2 μg/ml (cat# 07980, Stem Cell Technologies, Vancouver, Canada), EGF 20 ng/ml (Cat#315-09, Peprotech, London, UK) and FGF2 10 ng/ml (Cat#450-33, Peprotech, London, UK); penicillin 100 U/ml and streptomycin 150 μg/ml (15140-122, Gibco, Waltham, Massachusetts, USA). NSPCs were maintained for a maximum of 4 total passages to preserve a heterogeneous population and avoid cell selection issues. For proliferation/inhibition assays, 25.000 NSPC per well were seeded into 12-mm coverslips coated with laminin (L2020, Sigma-Aldrich, St Louis, MO, USA) as previously described (Silvestre et al., 2011), in order to avoid inhibition of cell proliferation by cell-to-cell contact. Cells were grown for 24h or 48h and a pulse of 10 μM bromodeoxyuridine (BrdU) (Cat#19-160, Sigma-Aldrich, St Louis, MO, USA) of 1h was given to detect cell proliferation. To assess either apoptotic and necrotic cell death pycnotic nuclei were identified with 4’,6-diamidino-2-phenylindole (DAPI) staining (Cat# D9542-5MG, Sigma-Aldrich, St Louis, MO, USA).

### 1.3 MTLE-HS model

For the stereotaxic injection of kainic acid (KA) (Cat#K0250, Sigma-Aldrich, St Louis, MO, USA) animals were anesthetized with a mixture of ketamine- (75mg/Kg; Ketamine, Bladel, The Netherlands) and medetomidine (1 mg/Kg; Sedastart; Pfizer, France) A single dose of the analgesic buprenorphine (1mg/kg) (Buprecare, Animalcare Lted) was administered subcutaneously. To induce status epilepticus 50 nL of a 20 mM solution of KA were delivered at the following coordinates: anteroposterior (AP), −1.9; lateral (L), −1.5; dorsoventral (DV), −2 mm; with bregma as reference. The delivery of the solution was performed using a micropump (Nanoject II, Drummond Scientific, Broomal, PA, USA) using a glass capillary as previously described (Sierra et al., 2015a). Control mice were given an intrahippocampal injection of 50 nL of 0.9% sterile sodium chloride (NaCl) (Cat#S5886, Sigma-Aldrich, St Louis, MO, USA). We confirmed the existence of initial seizures behaviorally using the Racine scale. All the animals included in the study reached category 4 or 5: clonic rearing and generalized tonic-clonic seizures with loss of righting reflex. All the MTLE mice reached at least one episode of category 4 o 5 convulsions during the period of 5-6h in which they were monitored the day of the surgery. The mortality rate is 10% with most of deaths occurring during or immediately after the surgery. The correct targeting of the injection was confirmed in the slices used for immunostaining.

### 1.4 Treatments

The stereotaxic injection of intrahippocampal zinc was performed using the coordinates described above using a 33G stainless steel cannula (outer diameter, 0.28 mm) connected to a Hamilton microsyringe (Hamilton, Bonaduz, Switzerland) as previously reported (Luzuriaga et al., 2019). The epidermal growth factor receptor (EGFR) irreversible inhibitor afatinib dimaleate (Ref. S7810, Selleck Chemicals LLC, Houston, TX, USA) was diluted with saline to a concentration of 2 mg/Kg and injected intrahippocampally immediately prior to KA delivery following the same steps as for the KA injection. In all conditions, the glass capillary or the cannula was left in the hippocampus for additional 5 min to avoid reflu. The skin of the skull was sutured and the mice were maintained in a thermal blanket until recovered from anesthesia. For zinc chelation N,N,N,N-Tetrakis (2-pyridylmethyl) ethylenediamine (TPEN) (Cat# P4413-50MG, Sigma-Aldrich, St Louis, MO, USA) solution was freshly prepared in 10%-ethanol (Cat#100983, Merck, Darmstadt, Germany) saline, and injected subcutaneously at 5 mg/Kg, twice a day as previously described(Kim et al., 2012). For intranasal treatment, 10 mg/Kg of gefitinib (Ref. S1025-SEL Selleck Chemicals LLC, Houston, TX, USA) were diluted with dimethyl sulfoxide (DMSO) (Cat#D8418, Sigma-Aldrich, St Louis, MO) plus 0.5% carboxymethylcellulose (Cat#419273, Sigma-Aldrich, St Louis, MO, USA) and 0.2% Tween 80 (Cat#P4780, Sigma-Aldrich, St Louis, MO, USA) following manufacturer instructions. Gefitinib administration started immediately after the stereotaxic KA injection, and continued twice a day for the first 3 days as previously described (Hanson et al., 2013; Pineda et al., 2013). BrdU (Cat#19-160, Sigma-Aldrich, St Louis, MO, USA) was diluted in sterile saline at 150 mg/kg concentration and administered through three intraperitoneal injections separated by 3h-intervals. BrdU injections were done 24h before sacrifice for the analysis at 3 dpKA and 3/7 days post administration of zinc (Cat#83265-250ML-F, Sigma-Aldrich, St Louis, MO, USA). To identify newborn cells at 14 dpKA, BrdU was given the last day of gefitinib treatment administration.

### 2.5 Real Time Quantitative Polymerase Chain Reaction

Ipsilateral hippocampi from WT mice were dissected at different time points (1.5h; 12h; 24h; 72h) post KA injection, immediately frozen with RLT buffer (Cat# 79216, Qiagen, Germantown, MD, USA) and processed for total RNA extraction using Micro RNeasy plus isolation kit (Cat#74034, Qiagen, Germantown, MD, USA). Reverse-transcription was done using a high-capacity reverse transcription kit (Cat#S7810, Thermo Fisher, Waltham, MA, USA). Real time quantitative polymerase chain reaction (RT-qPCR) was performed on a BioRad CFX96 device (BioRad, California, USA) using the following primers: hypoxanthine-guanine phosphoribosyltransferase (HPRT) forward 5’-GTT GGG CTT ACC TCA CTG CT −3’ reverse 5’-TCA TCG CTA ATC ACG ACG CT −3’; Glyceraldehydephosphate dehydrogenase (GAPDH) forward: 5’-CCA GTA TGA CTC CAC TCA CG −3’ reverse 5’ GAC TCC ACG ACA TAC TCA GC −3’; EGFR forward: 5’-GCC AAC TGT ACC TAT GGA TGT −3’ reverse 5’-GGC CCA GAG GAT TTG GAA GAA −3’; Fibroblastic growth factor receptor (FGFR) 1 forward: 5’ - CCA AAC CCT GTA GCT CCC TA −3’ reverse 5’ - TGA ACT TCA CCG TCT TGG CA −3’; FGFR2 forward: 5’-CCG AAT GAAGAC CAC GAC CA −3’ reverse 5’ TCG GCC GAA ACT GTT ACC TG −3’; Metallothionein (MTH) 1 forward: 5’ - TCA CCA CGA CTT CAA CGT CC −3’ reverse 5’ CAG TTG GGG TCC ATT CCG AG −3’; MTH2 forward: 5’-GCA TCT GCA AAG AGG CTT CC −3’ reverse 5’-AGT TGT GGA GAA CGA GTC AGG −3’; MTH3 forward: 5’ - GCT GCT GGA CTG GAT ATG GA −3’ reverse 5’ TTG CAT TTG TCC GAG CAG GT −3’. Each sample was normalized to endogenous GAPDH and to hypoxanthine guanine phosphoribosyltransferase (HPRT) that were used as housekeeper genes. Each reaction was performed at least twice in duplicates and the relative expression of each gene was calculated using the standard 2-ΔΔCt method (Livak and Schmittgen, 2001). HPRT and GAPDH were used as housekeeper genes.

### 1.5 Tissue and cell fixation and processing

Nestin-GFP mice or their WT (nestin-GFP-negative) control and treated animals were deeply anaesthetized with an intraperitoneal overdose of 2.5% of 2,2,2-Tribromoethanol (avertin) (Cat# T48402-25G, Sigma-Aldrich, St Louis, MO, USA) and were subjected to intra-cardiac perfusion with 30 ml PBS followed by 30 ml 4% paraformaldehyde (PFA) (Cat#158127, Sigma-Aldrich, St Louis, MO, USA) in 0.1 M PBS (pH 7.4). The brains were dissected and post-fixed for additional 3h at room temperature in 4% PFA and then rinsed with PBS. Serial sagittal vibratome sections were made (50μm-thick) using a Leica VT 1200S vibrating blade microtome (Leica Microsystems GmbH) and stored with PBS-0.02% sodium azide (Cat#S2002, Sigma-Aldrich, St Louis, MO, USA) at 4°C until use. The ipsilateral hemisphere was sliced sagittally in a lateral-to-medial direction, from the beginning of the lateral ventricle to the middle line, thus including the entire dentate gyrus (DG). The 50-μm slices were collected in 6 parallel sets, each set consisting of 12 slices, each slice 300 μm apart from the next. Cell cultures at the end of the treatments were fixed with PFA 4% dissolved with PBS-4% sucrose, rinsed with PBS and stored with PBS-0.02% sodium azide until use.

### 1.6 Immunofluorescence

For immunofluorescence, sections were incubated with blocking and permeabilization solution (PBS containing 0.25% Triton-X100 (Cat#93443, Sigma-Aldrich, St Louis, MO, USA) and 3% bovin serum albumin (BSA; Cat#A2153, Sigma-Aldrich, St Louis, MO, USA) for 3h at room temperature and then incubated overnight with the primary antibodies (diluted in the same solution) at 4°C. Cell cultures were permeabilized with PBS-0.3% triton-X100 1% BSA and then incubated with the respective primary antibodies overnight at 4°C. For BrdU and nestin staining, pretreatment with 2M hydrochlorhydric acid (HCl; Cat#1003141000, Sigma-Aldrich, St Louis, MO, USA) was applied during 20 minutes at 37°C followed by an immediate incubation with 0.1M tetraborate (Cat# 221732, Sigma-Aldrich, St Louis, MO, USA) for 10 minutes at room temperature before the blocking and permeabilization step. After the incubation, the primary antibody was removed and the sections or cell containing coverslips were washed with PBS three times for 10 minutes. Next, they were incubated with fluorochrome-conjugated secondary antibodies diluted in the permeabilization and blocking solution for 3h at room temperature. After washing with PBS, the sections or cell containing coverslips were mounted on gelatin coated slides with Fluorescent Mounting Medium (S3023, Dako Cytomation, Glostrup, Denmark). The following primary antibodies were used: GFP (1:1000, #GFP-1020) and Nestin (1:1000, #Nes) from Aves Laboratories (Tigard, USA); FGFR1 (1:200, #9740); HB-EGF (1:400, AF8239-SP, Abington, UK) from Cell Signaling Technologies (Massachussets, USA); EGFR (1:1000, ab52894) from Abcam, (Cambridge, UK); Ki67 (1:400) from Vector Laboratories (Burlingame, CA, USA); NeuN (1:200), Ki67 (1:1000) and glial fibrillary acidic protein (GFAP) (1:1500) from Dako Cytomation (Glostrup, Denmark); BrdU (1:300 Ab6326) from AbD Serotech (Kidlington, UK); and the following secondary antibodies: donkey anti-mouse, anti-rabbit, anti-goat or anti-rat Alexa −488, −568, −594, −647 or −680 secondary antibodies from Thermo Fisher (Waltham, USA); Anti Chicken secondary IgG Fluorescein (FITC; 603-702-C37, Tebu bio, Ile de France, France) DAPI at 1:1000 (Cat#D9542-5MG, Sigma-Aldrich, St. Louis, MO, USA) was used at counterstaining (for cell nuclei) when required.

### 1.7 Sholl analysis

3D-images containing single cells (at least n=50 per condition) were generated from z-stacks of saline+saline (n=4), saline+gefitinib (n=3), KA+saline (n=3), KA+gefitinb (n=5) Nestin-GFP mice using confocal microscopy. The cell profile was delimited using the “polygon” tool of the Image J and a mask was created adjusting threshold intensity to remove the background. Only complete cells including the cell body and cell processes were considered for the analysis. The voxel value was set up in 0.092 according to the resolution of the images. To determine cell geometry including the complex tree topology and the gross spatial arrangement we analyzed each condition using NeuronStudio 0.9.92 software (Computational Neurobiology and Imaging Center Mount Sinai School of Medicine New York, NY) (Rodriguez et al., 2006). Briefly, by defining the localization of the soma, the program automated the collection of the data for cell morphology including process length, cell volume and number of branch points and intersections. The results were exported into Excel spreadsheet. Graphic and statistical analysis comparing all four conditions were done using GraphPad Prism v5.

### 1.8 Western Blot

For western blot (WB) analyses cultured NSPCs or hippocampal tissue (at least n=3 per condition) were collected at indicated time-points for each experiment with RIPA buffer (Cat# A32963, Thermo Fisher, Waltham, USA) plus a protease inhibitor cocktail with the addition of phosphatase inhibitors (Cat#78441, Thermo Fisher, Waltham, USA). For cultured NSPCs, 400,000 cells per condition were used for cell signal transduction purposes. To study the effects of EGFR inhibition or zinc stimulation, cells were rinsed and cultured with starving media without growth factors during 2h to downregulate EGFR downstream signaling. Then, cells were stimulated either with 200 μM of zinc, 100 ng/mL of heparin-binding epidermal growth factor (HB-EGF) (SRP6050-10UG, Sigma-Aldrich, St Louis, MO, USA), 20 ng/mL of EGF and/or pre-treated with 2 μM gefitinib for 1h. After homogenization, samples were centrifuged at 14,000 rpm for 10 min. Supernatant protein (20 μg) was loaded in a 10%-Tris-Glycine gels and transferred to a nitrocellulose membrane (Cat#88018 Life technologies, Carlsbad, CA, USA). Blots were blocked in 5%, nonfat, dry milk in TBS-T (150mM NaCl, 20 mM Tris-HCl, pH 7.5, 0.05% Tween 20) and then incubated with the following antibodies: Phospho-specific antibodies against EGFR (Tyr1068, 1:1000, #3777, Tyr845, 1:1000, #2231), AKT (Ser473, 1:1000, #9271), ERK (Thr202/Tyr204, 1:2000, #4370) and STAT3 (Tyr705) (1:1000, #9131, Cell Signaling, Massachusetts, USA) and total EGFR (1:1000, ab52894, Abcam, Cambridge, UK), AKT (1:1000, #9272), ERK (1:1000 #4695), STAT3 (1:2000, #4904) and FGFR (1:1000, #9740) from Cell Signaling (Massachusetts, USA) and MTH1/2 (1:1000, MA1-25479, Thermo Fisher, Waltham, USA). Anti-ß-actin (1:1000, #3700, Cell Signaling, Massachusetts, USA) was used for loading control and Ponceau staining (P7170, Sigma-Aldrich, St Louis, MO, USA) was carried out to check protein and band migration. After three washes in TBS-T, blots were incubated with 1:1000 of either anti-mouse or anti-rabbit IgG HRP-conjugated (Life technologies, Carlsbad, USA) and developed by enhanced chemiluminescent (ECL) SuperSignal (#34095, Thermo Fisher, Waltham, USA) WB analysis system. ECL signal was captured using a ChemiDocTM Imaging System (Bio-Rad, California USA). Quantification of the signal was performed by densitometric scanning of the membrane using GelPRO analyzer software (Media Cybernetics 1993-97).

### 1.9 Enzyme Linked Immunosorbent Assay

HB-EGF content was determined in duplicates by the R&D DuoSet Immunoassay (R&D) as described previously (Canals et al., 2004). Briefly, either cellular pellet or brain tissue was sonicated using a buffer containing Hepes 25 mM (Cat#54457, Sigma-Aldrich, St Louis, MO, USA), MgCl2 5 mM (Cat# M4880, Sigma-Aldrich, St Louis, MO, USA) ethylene glycol-bis(β-aminoethyl ether)-N,N,N’,N’-tetraacetic acid (EGTA; Cat#E3889, Sigma-Aldrich, St Louis, MO, USA) 1 mM, ethylenediaminetetraacetic acid (EDTA; Cat#E6758, Sigma-Aldrich, St Louis, MO, USA) 1.3 mM, phenylmethyl sulfonide fluoride (PMSF; Cat# PMSF-RO, Sigma-Aldrich, St Louis, MO, USA) 1mM and a protease and phosphatase cocktail of inhibitors (Thermo Fisher, Ref. 88665 and 78420 respectively). For cultured NSPCs, 250,000 cells were seeded in a 96 well plate with 150 μL of culture media without growth factors and the cell containing media was collected after 72h of plating. We analyzed 100 μL of both the supernatant and lysed cell pellet from the NSPCs culture, diluting them 1:1 with reagent buffer 1X. In parallel conditions, right after 72h of culture, the number of cells was counted using a Bio-Rad TC20 automated cell counter (Bio-Rad Laboratories, California, USA), using trypan blue (Cat#302643, Sigma-Aldrich, St Louis, MO, USA) to mark and exclude death cells. Next, we determined the amount of HB-EGF per cell through enzyme linked immunosorbent assay (ELISA; Cat# DY8239-05, R&D Systems, Minneapolis, MN, USA). For analysis of HB-EGF levels in brain tissue, mice were deeply anesthetized in a CO2 chamber at 1.5h, 12h, 24h and 72h post-KA or saline administration (n=4). Their hippocampi were removed and the DG was dissected out on ice and rapidly frozen using CO2 pellets. Samples were homogenized in the abovementioned lysis buffer and centrifuged 20 min at 14,000 rpm at 4°C. Supernatants were collected and the total protein content was analyzed using a basic colorimetric assay (BCA) protein assay kit (23227, Pierce, Dallas, USA). One hundred μg of protein were loaded for each time point and diluted 1:1 in Reagent diluent. Both *in vitro* and *in vivo* quantifications were performed using the following four-parameter logistic (4-PL) curve fit: y=d+(a-d)/(1+(x/c)^b), where x and y were the independent and dependent variables respectively, a and d were the minimum and maximum values measured, c was the point of inflection (halfway between a and d and b was hill’s slope of the curve (steepness of the curve at point c). For in vitro values were normalized and expressed as picogram of HB-EGF per mL and for in vivo values were calculated as pg of HB-EGF per μg of tissue protein.

### 2.10 Danscher staining

The indirect autometallographic Danscher method is based on the precipitation of metal ions as insoluble salts that can be visualized by microscopy. Twenty mg/Kg of sodium selenite (S5261-10G, Sigma-Aldrich, St. Louis, USA) were injected intraperitoneally in n=3 control and n=3 KA Nestin-GFP animals 30 minutes before perfusion with a PFA 4 %-Glutaraldehyde 0.5 % (Cat#16536-05, Electron Microscopy Sciences, Hatfield, PA, USA) mixture. Brains were dissected and serial vibratome sections were made (50 μm-thick) using a Leica VT 1200S vibrating blade microtome (Leica Microsystems GmbH). Autometallography of zinc-selenium nanocrystals was developed as previously described (López-García et al., 2002). Granule size and quantification was carried out determining the region of interest (ROI), in this case GFP-expressing cells (15-20 cells per condition) and using the particle analyzer of ImageJ software.

### 2.11 Image capture

Immunofluorescence images were collected employing the 20X or the 40X oil-immersion objective of a Leica SP8 (Leica, Wetzlar, Germany) laser scanning microscope and their corresponding manufacturer’s software (LAS X, Leica Microsystems, Wetzlar, Germany). For the quantification of total areas of the DG the 10X objective was used to completely visualize the DG in each section. The zinc granules in Figure 4 were quantified using an Olympus BX31 microscope with an attached Olympus DP72 high sensitivity camera (Olympus, Shinjuku, Japan). The brightfield channel was used for zinc granules and was overlapped with the green fluorescence channel was used for Nestin-GFP signal. The signal from each fluorochrome was collected sequentially, and controls with sections stained with the fluorochrome was performed to confirm the absence of signal leaking into different channel. Brightness, contrast, and background were adjusted equally for the entire image without any further modification. Al images shown are flat projections from z-stacks ranging from 10 (typically for individual cell images) to 20 μm of thickness. 4-5 z-stacks located at random positions in the DG were collected per hippocampal section, and 4-6 sections per series were analyzed, depending on the experiment. For cultured NSPCs, 5 pictures of 4 μm-thick random z-stacks of the coverslips were collected per sample and condition.

### 3.3 Cell quantification

Quantitative analysis of cell populations *in vivo* was performed by design-based (assumption free, unbiased) stereology using a modified optical fractionator sampling scheme as previously described (Encinas et al., 2006, 2011; Encinas and Enikolopov, 2008). For cell densities, quantifications were done maintaining the same z-stack size and settings among conditions. The values were normalized to the volume of the SGZ+GCL (subgranular zone+ granule cell layer). For total numbers, the whole area of the DG was determined in every slice of a series and then multiplied by the thickness of the GCL, that was measured at least in three different points in each brain slice to obtain an average. The obtained value was multiplied by the number of series obtained during vibratome sectioning (5 or 6) to obtain the volume of the whole DG.

NSCs were counted following previously described criteria for their identification (Kronenberg et al., 2003; Encinas and Enikolopov, 2008), defining them as radial glia-like cells positive for Nestin-GFP and GFAP with the soma located in the SGZ or the lower third of the GCL and with a process extending from the SGZ towards the molecular layer through the GCL. Amplifying neural progenitors (ANPs) were defined as Nestin-GFP-positive cells devoid of GFAP immunostaining and with none or short horizontal processes. The cell cycle marker Ki67 or administration of BrdU and its posterior immunolabeling were used to identify proliferating cells. Proliferation of cultured NSPCs was measured by the incorporation of BrdU. The relative proportions of BrdU-positive NSPCs were referred to the total number of cells (DAPI staining) quantified per coverslip and conditions. The proliferation in all the conditions was referred to the basal proliferation of the control condition.

To measure cell death, apoptotic o necrotic cells were defined as cells with small condensed nuclei with abnormal morphology (pyknotic/karyorrhectic). GCD (distance from the SGZ to the molecular layer) was measured in DAPI- or NeuN-stained sections and the thickness of the GCL was measured at least in three points in each slice.

For the analysis of EGFR, an automated batch analysis for ImageJ was used. In brief, the area was measured as the pixels occupied by EGFR. The ROI was selected in the Nestin-GFP channel using the tool “Color threshold, HSB color space” to adjust the ROI for NSCs. The generated mask was overlayed as the ROI in the rest of channels. Automated quantification was obtained with the “find maxima” option, using the same “noise-tolerance” for all conditions, obtaining the total area occupied by EGFR. For the analysis of EGFR expression in cell culture, at least 25 mitotic or interphasic cells were analyzed. Using the tool “polygon” of the ImageJ the outline of the cells in brightfield channel was delimited to determine cell area and create the respective ROI for each cell, representing cell area and the ratio of pixel intensity of EGFR signal per cell area.

We analyzed the whole dentate gyrus for absolute numbers of cells (ki67, dead cells…) and the medial half of the hippocampus for analyses related to NSCs and React-NSCs. As the dentate gyrus changes its orientation so do NSCs and is more difficult to identify them in the most lateral aspect of the hippocampus.

### 3.4 Statistical analysis

The statistical testing was implemented using Sigma-Aldrich Plot version 12 (Systat Software, Chicago, USA), except from the Sholl analysis in Figure 3F, 3G and 3H in which GraphPad Prism version 5 (San Diego, CA, USA) was used. Comparisons between multiple groups were made using one-way analysis of variance (ANOVA) or Kruskal-Wallis when the data did not comply with normality. The analyses were followed by multiple comparisons; All pairwise in Figures 3E-H, 5C-K and Supplemental Figures 3G, 4F and 5C; and versus the control group in Figures 1B-G and Supplemental Figures 4A-D. The Holm-Sidak post hoc method was used for multiple comparisons after one-way ANOVA in all cases except for Figure 5F, where the Fisher LSD method was performed. Kruskal-Wallis was followed by the post hoc Dunnet’s method in Figures 1E-F and Supplemental Figure 4A for MT1; the post hoc Dunn’s method in Supplemental Figure 4D-F; and by the post hoc Student-Newman-Keuls method in Figure 5C and 5H and Supplemental Figures 3B, 3G, and 5C. Comparisons between only two groups were made using two-tailed unpaired Student’s t-test or Mann Whitney U test when the data did not comply with normality assumption. All statistical tests and the number of independent experiments can be found in the figure legends. p<0.05 was considered as statistically significant. Results were presented as mean ± standard error mean (SEM).

